# Coupling a Live Cell Directed Evolution Assay with Coevolutionary Landscapes to Engineer an Improved Fluorescent Rhodopsin Chloride Sensor

**DOI:** 10.1101/2022.01.20.474067

**Authors:** Hsichuan Chi, Qin Zhou, Jasmine N. Tutol, Shelby M. Phelps, Jessica Lee, Paarth Kapadia, Faruck Morcos, Sheel C. Dodani

**Author notes:** H.C., Q.Z., J.N.T., and S.M.P. contributed equally to this work.

## Abstract

Our understanding of chloride in biology can be accelerated through the application of fluorescent protein-based sensors in living cells. Laboratory-guided evolution can be used to diversify and identify sensors with specific properties. Recently, we established that the fluorescent proton-pumping rhodopsin *wt*GR from *Gloeobacter violaceus* can be converted into a fluorescent sensor for chloride. To unlock this non-natural function, a single point mutation at the Schiff counterion position from D121V was introduced into *wt*GR fused with cyan fluorescent protein (CFP) resulting in GR1-CFP. Here, we have integrated coevolutionary analysis with directed evolution to understand how the rhodopsin sequence space can be explored and engineered to improve this starting point. We first show how evolutionary couplings are predictive of functional sites in the rhodopsin family and how a fitness metric based on sequence can be used to quantify known proton-pumping activities of GR-CFP variants. Then, we couple this ability to predict potential functional outcomes with a screening and selection assay in live *Escherichia coli* to reduce the mutational search space of five residues along the proton-pumping pathway in GR1-CFP. This iterative selection process results in GR2-CFP with four additional mutations, E132K, A84K, T125C, and V245I. Finally, bulk, and single fluorescence measurements in live *E. coli* reveal that GR2-CFP is a reversible, ratiometric fluorescent sensor for extracellular chloride with an improved dynamic range. We anticipate that our framework will be applicable to other systems, providing a more efficient methodology to engineer fluorescent-protein based sensors with desired properties.

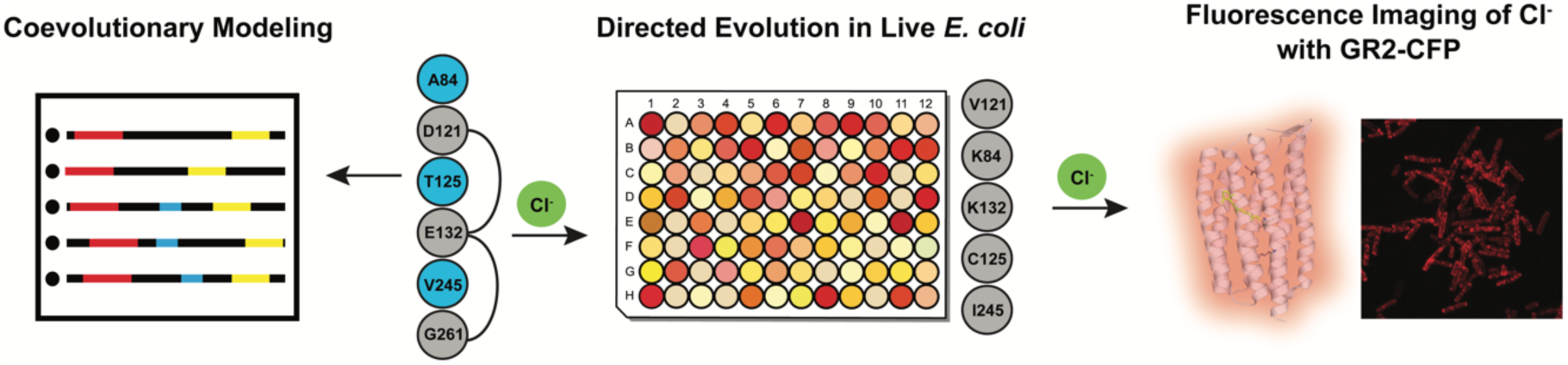

## Introduction

Chloride is universally a vital ion for life.^1–3^ Unicellular and multicellular organisms alike have evolved membrane-bound transporters to mobilize chloride for a wide range of biological functions.^1,3–5^ Our ability to visualize and intercept these dynamic processes has been enabled by protein-based fluorescent sensors for chloride.^5–8^ To date, these have in large part been derived from the green fluorescent protein (GFP) found in the jellyfish *Aequorea victoria*.^9^ Early efforts to engineer GFP led to the discovery of chloride-sensitive variants, YFP, E^1^GFP, and Topaz.^10–14^ These three starting points have been extensively engineered, providing insights into how the sequence of GFP is connected to this unique function. Protein engineering workflows have primarily relied on site-directed/saturation mutagenesis coupled to selection in *Escherichia coli* extracts or cell-free systems.^5,6,8,13–18^ As a complementary strategy, we have used bioinformatic and structure-guided approaches to identify chloride-sensitive fluorescent proteins with new properties, including the naturally occurring phiYFP from the jellyfish *Phialidium sp. SL-2003* and the engineered monomer mNeonGreen from the cephalochordate *Branchiostoma lanceolatum*.^19,20^

Expanding from the GFP family (Pfam: PF01353), we recently showcased a *de novo* design strategy to confer chloride-sensitivity into a fluorescent, outward proton-pumping rhodopsin from the cyanobacterium *Gloeobacter violaceus* (*wt*GR).^21^ Like other microbial rhodopsins (Pfam: PF01036), *wt*GR is a membrane-spanning protein with a visible light-harvesting Schiff base chromophore (SBC) formed from the condensation of all-*trans*-retinal and a lysine residue (K257 in *wt*GR).^22–25^ Outside its natural context, the SBC can also absorb near-infrared light to access a fluorescent state.^23,26–31^ Since the aspartate counterion (D121 in *wt*GR) can stabilize and tune the spectroscopic properties of the SBC, we hypothesized that a chloride ion could do the same.^21^ To engineer this non-natural function, we first developed a high-throughput protein engineering workflow in live *E. coli*. A fusion between *wt*GR and cyan fluorescent protein (CFP) was used to generate a chloride-insensitive starting point with a red-shifted, ratiometric fluorescent output. Site-saturation mutagenesis at D121 resulted in the identification of GR1-CFP where mutation to valine not only resulted in a turn-on fluorescence response to 400 mM chloride (ΔF_GR_/F_CFP_ ≈ 2.0 at pH 5) but also eliminated pumping activity in live *E. coli*.^21^

Even though GR1-CFP does not pump protons nor chloride ions, the residues that define the parent proton pump are still present. In *wt*GR, the proton from the SBC is first donated to the aspartate counterion (D121), which is stabilized through hydrogen bonding interactions with T125 and released on the extracellular side.^25,32,33^ Then, the glutamate donor (E132) can re-protonate the SBC on the cytoplasmic side.^25,32,33^ This DTE motif is a characteristic sequence that defines a proton-pumping rhodopsin where the glutamate can be substituted with aspartate as found in the well-studied bacteriorhodopsin (BR) from the archaeon *Halobacterium salinarum*.^25,29,34–36^ Along the proton-pumping pathway, residues A84 and G261 in *wt*GR are conserved in BR and related proton pumps, but the terminal proton acceptor position is substituted with V245 (Figure S1).^25,35^ Since GR1-CFP has the D121V mutation, the context of its interactions with these and other residues that are essential for the parent function has changed. To better understand this and how the sensor capabilities of GR1-CFP could be improved through protein engineering, here, we have integrated coevolutionary analysis with our live cell directed evolution assay.

As proteins evolve to preserve or acquire new functions, their change in amino acid composition leaves a pattern that can be useful to identify critical sites relevant for function. Although amino acid conservation has been of use for many years, patterns of covariation can also provide important clues about epistatic interactions relevant to folding, catalytic activity, and interactions. Statistical modeling of coevolving sites at the amino acid level has become a useful technique to study structure, function, and interactions of biomolecules. An efficient and accurate methodology to study residue coevolution in protein families is Direct Coupling Analysis (DCA).^37,38^ DCA is a global statistical model that reliably captures evolutionary constraints on amino acid sites in multiple sequence alignments (MSAs) of biomolecular families. Coevolutionary information has been successfully used in diverse applications, including the determination of protein 3D structures,^39,40^ discovery of protein conformational dynamics,^41^ prediction of protein interaction partners and their specificities,^42–44^ protein-protein interaction interface inference,^45–49^ mutational landscapes for protein–nucleic acid recognition,^50^ as well as functional protein design.^51,52^ To our knowledge, this approach has yet to be explored for fluorescent protein-based sensors. In this study, we investigate how the use of evolutionary information and the determination of a sequence mutational landscape in the rhodopsin family provides a unique strategy to engineer GR1-CFP with potential extensions to fluorescent protein-based sensors in general. By integrating coevolutionary modeling, mutagenesis, screening, and an iterative selection strategy (Figure 1), we can effectively search the functional mutational space of the residues A84, T125, E132, V245, and G261 in GR1-CFP to generate the improved variant GR2-CFP.

**Figure 1.**
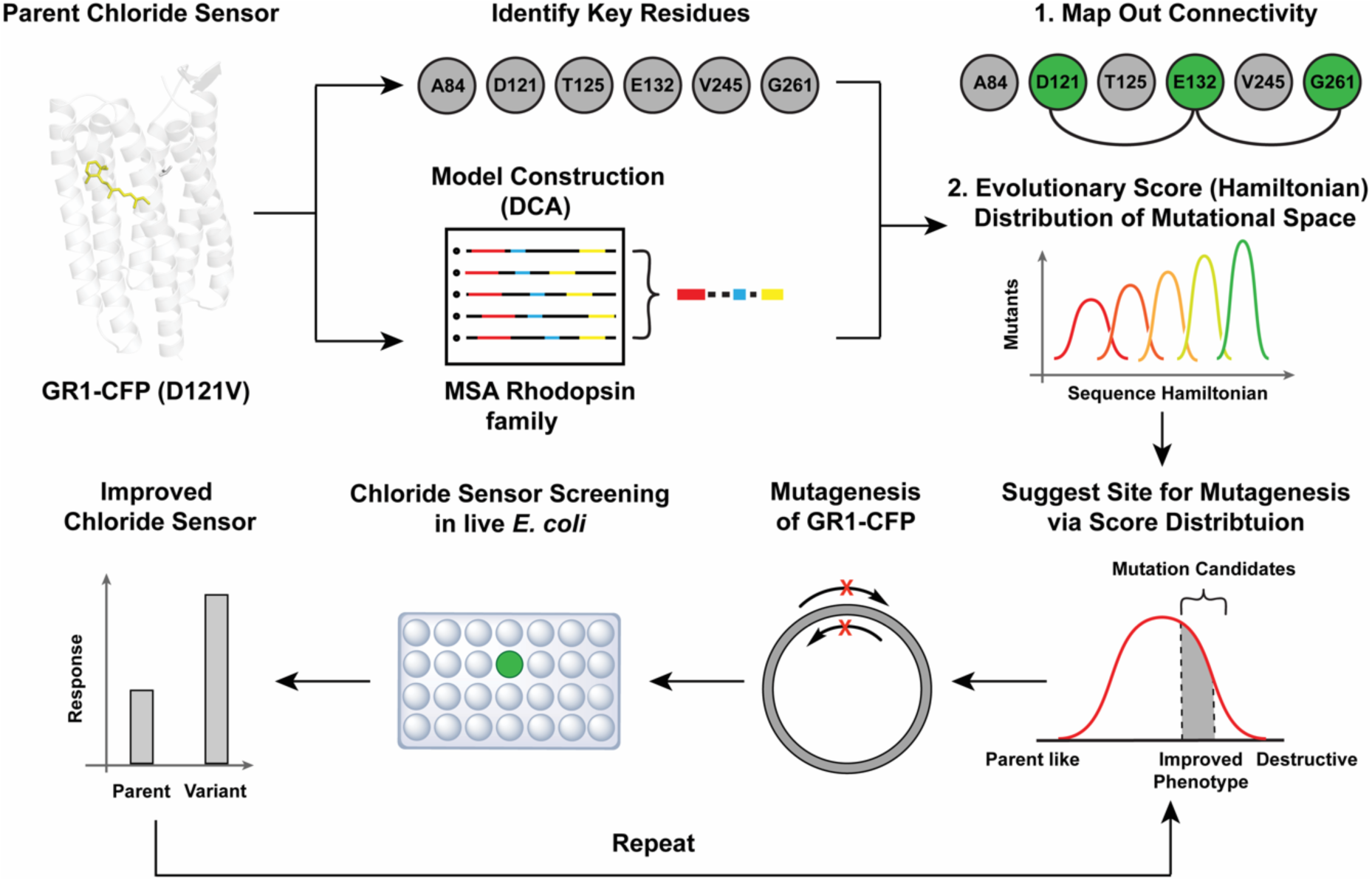
The workflow to couple directed evolution and coevolutionary landscapes to improve the properties of the fluorescent rhodopsin chloride biosensor GR1-CFP was divided into three steps. First, residues along the proton-pumping pathway of wild-type *Gloeobacter violaceus* rhodopsin (*wt*GR) were identified from the literature and projected onto a direct coupling analysis of the rhodopsin family to build a connectivity map. Then, the available mutational space was refined based on the Hamiltonian distribution to guide the mutagenesis order. In parallel, an iterative site-saturation mutagenesis, screening, and selection strategy in live *E. coli* was developed to identify variants with an improved turn-on fluorescence response to chloride.

## Results

### Coevolutionary model of the rhodopsin family

We first used Direct Coupling Analysis to capture the coevolutionary information from the rhodopsin family sequences with two types of metrics: i) direct information (DI) that quantifies the degree of coupling or coevolution among two residue positions in the MSA of the rhodopsin family and ii) Hamiltonian scores that provide a global measure related to the probability of a given sequence to be a member of the family (Figure 1, step 2, Figure S2). DI pairs were used to identify the key coevolutionary pairwise correlations between residues in *wt*GR. A Hamiltonian score is calculated as a sum of all the pairwise and single site parameters, in the joint probability distribution of a family (see Methods). Since members of the rhodopsin family are light-activated ion pumps and not known to be fluorescent sensors for chloride, the Hamiltonian scores were used to select mutants whose parameters are not necessarily optimized for family likeness.

### Identification of evolutionary coupled residues involved in the proton-pumping pathway of wtGR

In this model, we focused on position 121 and five residues along the proton-pumping pathway of *wt*GR described above (Figure 2A). We performed DCA for the protein sequences in the rhodopsin family to quantitatively investigate these residues and validate our modeling in identifying relevant interacting amino acids (see Methods). First, we generated a connectivity network based on the top 300 DI pairs (Figure S2, Supplemental File 1). The specific DI pairs that are related to the proton-pumping pathway residues are shown as a network in Figure 2B. We identified strong direct evolutionary interactions among 84, 125, 121, and 132. Some of these residue pairs are close in the GR1 homology model that is based on the crystal structure of *wt*GR (PDB ID: 6NWD).^24^ For instance, the DI pairs 84-121 and 84-125 may form physical contacts relevant for folding and function, as the contact distances are 5.2 Å and 4.7 Å, respectively (Table S1). For the 121-132 pair, the distance is 11.3 Å, which is in line with the fact that they are the proton donor/acceptor pair for the SBC. In addition to the coevolutionary interactions to the selected proton-pumping pathway residues, we found other residues that are highly coupled, suggesting additional relevant interactions that are essential for the natural function of *wt*GR. For example, residues 132 and 77 are key residues that allow for inward directed diffusion of protons under certain conditions.^33^ The strong interaction between residues 121 and 87 is consistent with how these residues form a salt-bridge to functionally position residues for the light-induced proton transfer.^53,54^ Interestingly, by labeling these residues in the structural model, we observe that they form a mesh that surrounds the proton-pumping pathway (Figure S3). The other two residues, 245 and 261, do not directly interact with the other selected proton-pumping pathway residues but are connected through intermediate residues. For example, both residues 261 and 84 are coupled to a single residue 129. This interaction is also reflected in the distance measurements. The 84-261 pair is far apart (∼12 Å), while the 84-129 and 129-261 pairs are closer (8.4 Å and 6.9 Å, respectively). The latter pairs could potentially form physical contacts required for folding and function (Table S1). Other residues from our model such as 87, 88, 253, and 257 are known to be near the SBC and essential for the native proton release complex.^25^ This analysis supports the idea that i) direct couplings do capture physical interactions relevant to the parent proton pumping function and ii) evolutionary couplings can in principle be used to study changes that can affect functional phenotypes.

**Figure 2.**
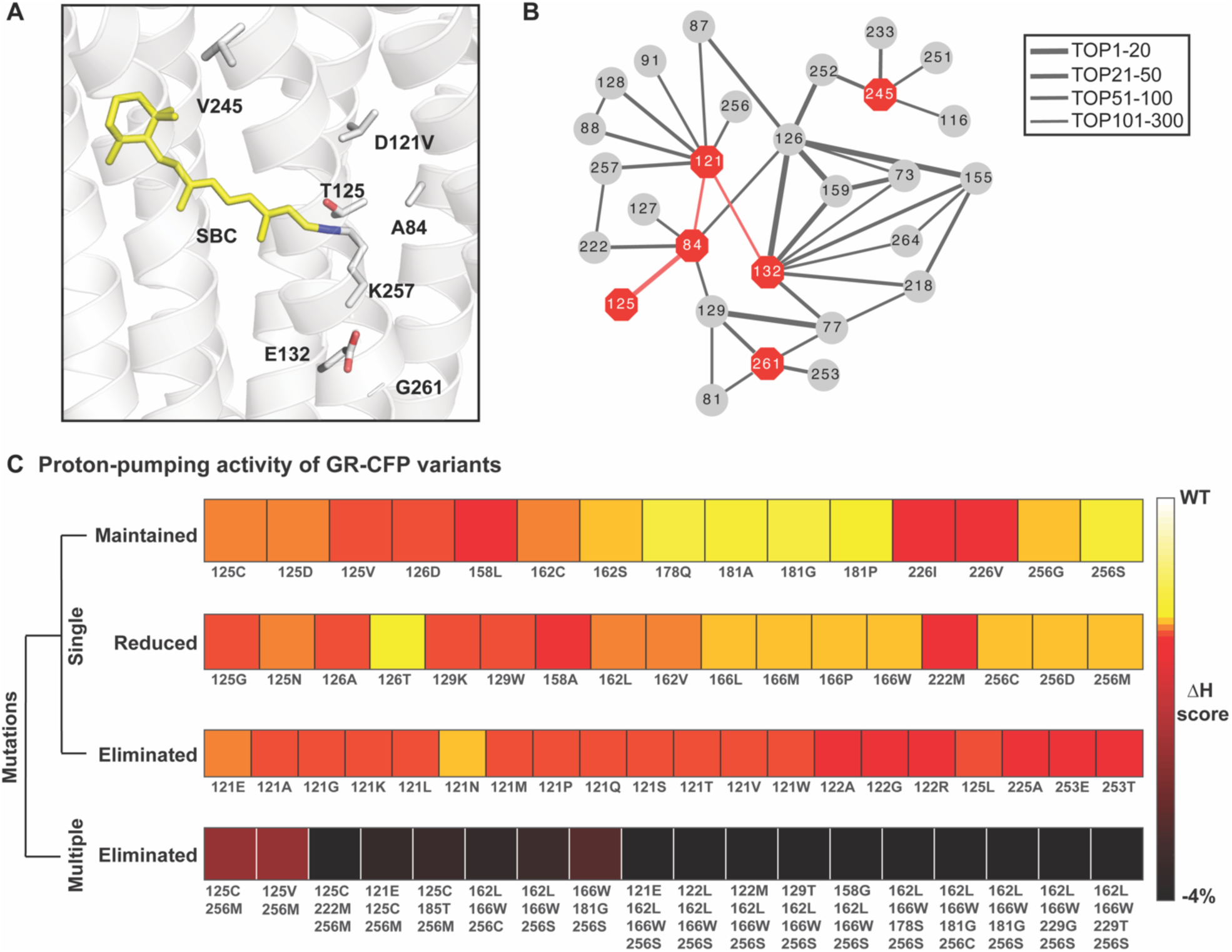
Coevolutionary information reveals important functional residues in the rhodopsin family. (A) Homology model of the rhodopsin chloride biosensor GR1 engineered from the proton-pumping rhodopsin *Gloeobacter violaceus* (*wt*GR, PDB ID: 6NWD)^24^ without CFP. The residues along the proton-pumping pathway selected for mutagenesis are shown. All residues are shown as gray sticks with the oxygen atoms in red and the nitrogen atom in blue each labeled with the single letter amino acid code and position number. The Schiff base chromophore (SBC) formed from all-*trans*-retinal (yellow sticks) and K257 is shown. (B) Connectivity map of the coupled pairs for the selected residues in the top 300 direction information (DI) pairs (Figure S2, Supplemental File 1). The position numbers of the residues along the proton-pumping pathway are shown as red octagons and the evolutionary coupled residues as gray circles. Each pair of coupled residues is connected with a line. The thickness of the lines correlates with a higher rank of the DI pair indicating a stronger evolutionary coupling strength. (C) Comparison between proton-pumping activity levels and the Hamiltonian scores for previously reported GR-CFP variants to validate the coevolutionary method.^23^ The variants with the same level of proton-pumping activities are grouped together in the same row. The relative changes of the Hamiltonian scores are illustrated in the heatmap from white (corresponding to a *wt*GR phenotype) to black with each mutation listed with the single letter amino acid code and position number.

### Coevolutionary modelling is predictive of proton-pumping activities of wtGR-CFP and variants

To test the accuracy of the coevolutionary model in predicting functional outcomes in the rhodopsin family, we first studied the effect of mutations in previously reported variants of *wt*GR-CFP.^23^ For each variant, we correlated the mutations and proton-pumping activities (maintained, reduced, or eliminated) with the calculated Hamiltonian score. These results are illustrated in the heatmap of Figure 2C. With respect to the wild-type, we observed a gradual decrease in the relative Hamiltonian scores for variants that have no proton-pumping activity. With this information at hand, we created a Hamiltonian-based classifier that predicts if a variant will eliminate proton-pumping activity. We found that such classification task can be done with high accuracy (Area Under the Curve, AUC = 0.90; Figure S4) showcasing the predictive power of coevolutionary information for functional features in the rhodopsin family.

### Iterative site-saturation mutagenesis of GR1-CFP guided by the Hamiltonian score

Having established the predictive power of the DI and Hamiltonian metrics, we utilized this methodology to guide the mutagenesis of GR1-CFP. To achieve this, we calculated i) the Hamiltonian score for every possible variant at position 121 and five residues selected along the proton-pumping pathway and ii) the mean of the relative Hamiltonian scores changes for all the mutations in the same position (see Methods) to illustrate the effect of changes at each site (Figure 3). Position 121 showed the most negative relative change among all the six residues. This suggested that to increase the fluorescence response to chloride, we should select positions with more negative relative changes for mutagenesis, thus deviating from the parent function.

**Figure 3.**
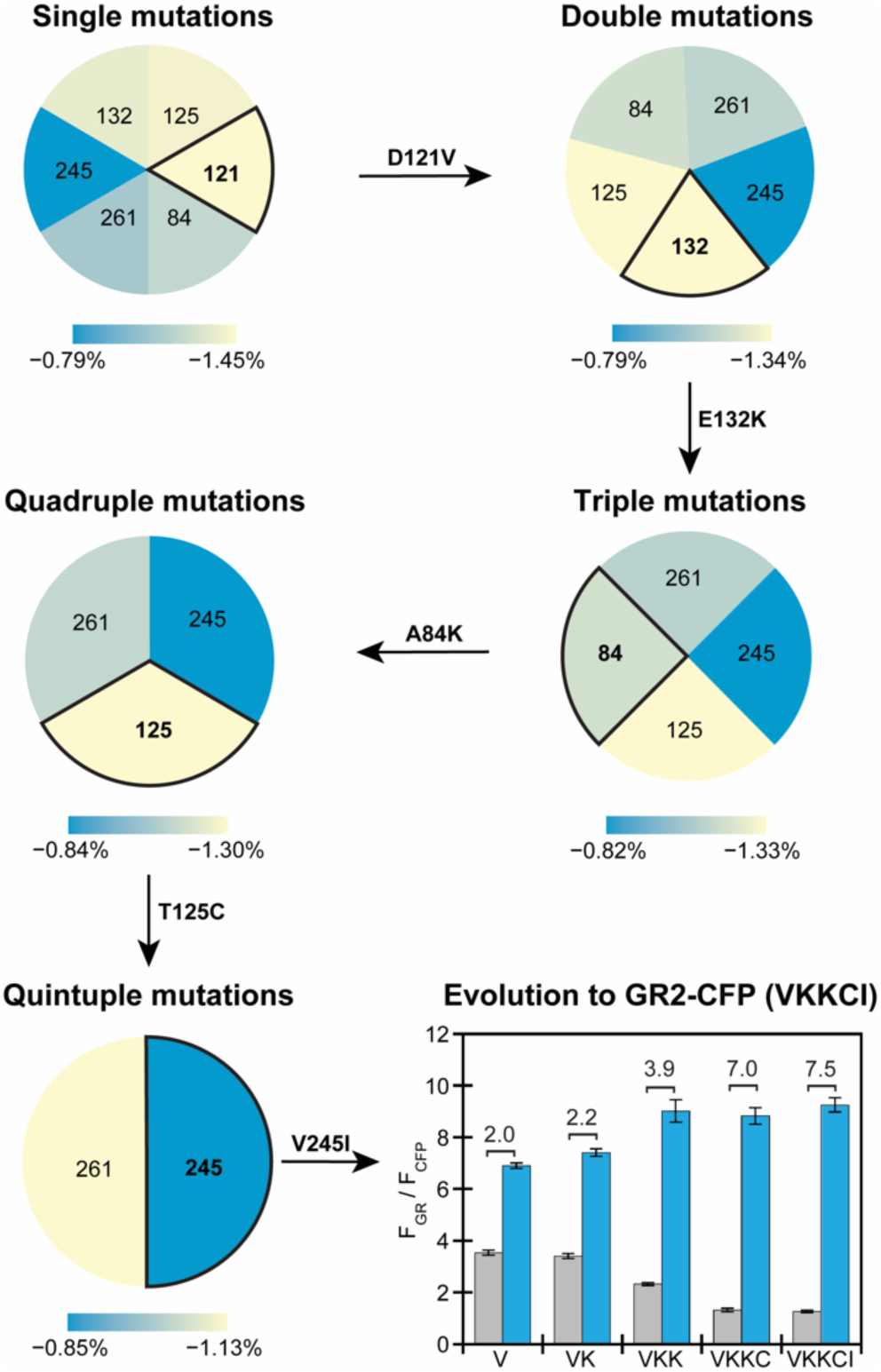
Iterative site-saturation mutagenesis of residues along the proton-pumping pathway in GR1-CFP guided by a Hamiltonian score prediction results in GR2-CFP with an improved turn-on fluorescence response. From single mutations to quintuple mutations, the mutational space was refined by comparing the relative changes of the average Hamiltonian score from blue to yellow (largest change) between all the possible mutations at the positions listed with respect to the parent. In each step, the position with the largest change in the Hamiltonian score was selected for mutation. The identity of the variant with an improved fluorescence response to chloride is shown above the arrow as the starting parent for the next site targeted for mutagenesis. The turn-on fluorescence response of GR1-CFP and each improved variant generated from site-saturation mutagenesis and screening in live *E. coli* is summarized in the bar graph. Each bar represents the ratio of the normalized integrated emission (F_GR_/F_CFP_) in the absence (gray) and presence (blue) 400 mM sodium chloride in 50 mM sodium acetate at pH 5. For each bar, the average of twelve biological replicates from three technical replicates with standard error of the mean is shown. The turn-on response to chloride (ΔF_GR_/F_CFP_) is shown for each variant. For the rhodopsin, excitation was provided at 570 nm and emission was collected and integrated from 630–760 nm (F_GR_). For CFP, excitation was provided at 390 nm and emission was collected from 435–520 nm (F_CFP_). Emission spectra are shown in Figure S7, Figure S8.

In our experimental approach, site-saturation mutagenesis was used to generate all possible amino acid substitutions at each position with the largest Hamiltonian score change (Figure 3, Table S2, and Figure S5–S8). We first fixed position 121 as valine, and selected position 132 for mutagenesis. This residue had the largest score changes and co-evolved with many proton-pumping related residues (Figure 2B, Figure S2). The library was screened in live *E. coli* and variants with an improved turn-on, ratiometric response to 400 mM sodium chloride versus GR1-CFP (ΔF_GR_/F_CFP_ ≈ 2.0 at pH 5) were rescreened prior to Sanger sequencing to identify the mutations. This initial screen generated GR1-E132K-CFP (ΔF_GR_/F_CFP_ ≈ 2.2 at pH 5), which served as the new starting point. Next, position 125 was selected, but no improved variants were experimentally identified. We proceeded to position 84 that had the next largest score change and identified GR1-E132K-A84K-CFP (ΔF_GR_/F_CFP_ ≈ 3.9 at pH 5). Continuing this cycle of modeling selection, mutagenesis, and screening, we identified the final variant with four mutations GR1-E132K-A84K-T125C-V245I-CFP or GR2-CFP (ΔF_GR_/F_CFP_ ≈ 7.5 at pH 5). No improved variants were identified at position 261.

### Characterization of GR2-CFP in live E. coli cells using bulk and single cell fluorescence measurements

To understand how the accumulated mutations altered sensing properties, bulk fluorescence measurements were directly carried out in live *E. coli* cells expressing GR2-CFP (Figure S9). In these experiments, the emission of GR2 was normalized to the emission of CFP, thus providing a ratiometric output for exogenous chloride supplementation. At pH 5, excitation of apo GR2 at 570 nm results in no fluorescence signal above the background of the cells alone, but excitation of CFP at 390 nm results in an emission maximum centered at 485 nm, confirming the expression of GR2-CFP (Figure 4A, Figure S10–S13). However, in the presence of up to 400 mM sodium chloride, GR2-CFP has a broad emission centered at 705 nm with a ΔF_GR_/F_CFP_ = 7.5 ± 1.5. This turn-on response arises from GR2 as the fluorescence intensity of CFP remains unchanged. Fitting of the titration data to a single site binding model results in an apparent dissociation constant (*K*_d_) for chloride binding to GR2-CFP of 53 ± 14 mM. We next evaluated if GR2-CFP could operate at more physiologically relevant pH (Figure S14, Figure S15). In the absence of chloride, GR2-CFP also has no measurable fluorescence at pH 6 and 7. However, surprisingly, the fluorescence ratio increases (ΔF_GR_/F_CFP_ = 3.8 ± 0.35) with up to 400 mM sodium chloride at pH 6 (*K*_d_ = 731 ± 31 mM). No fluorescence response is observed at pH 7 (Figure S15).

**Figure 4.**
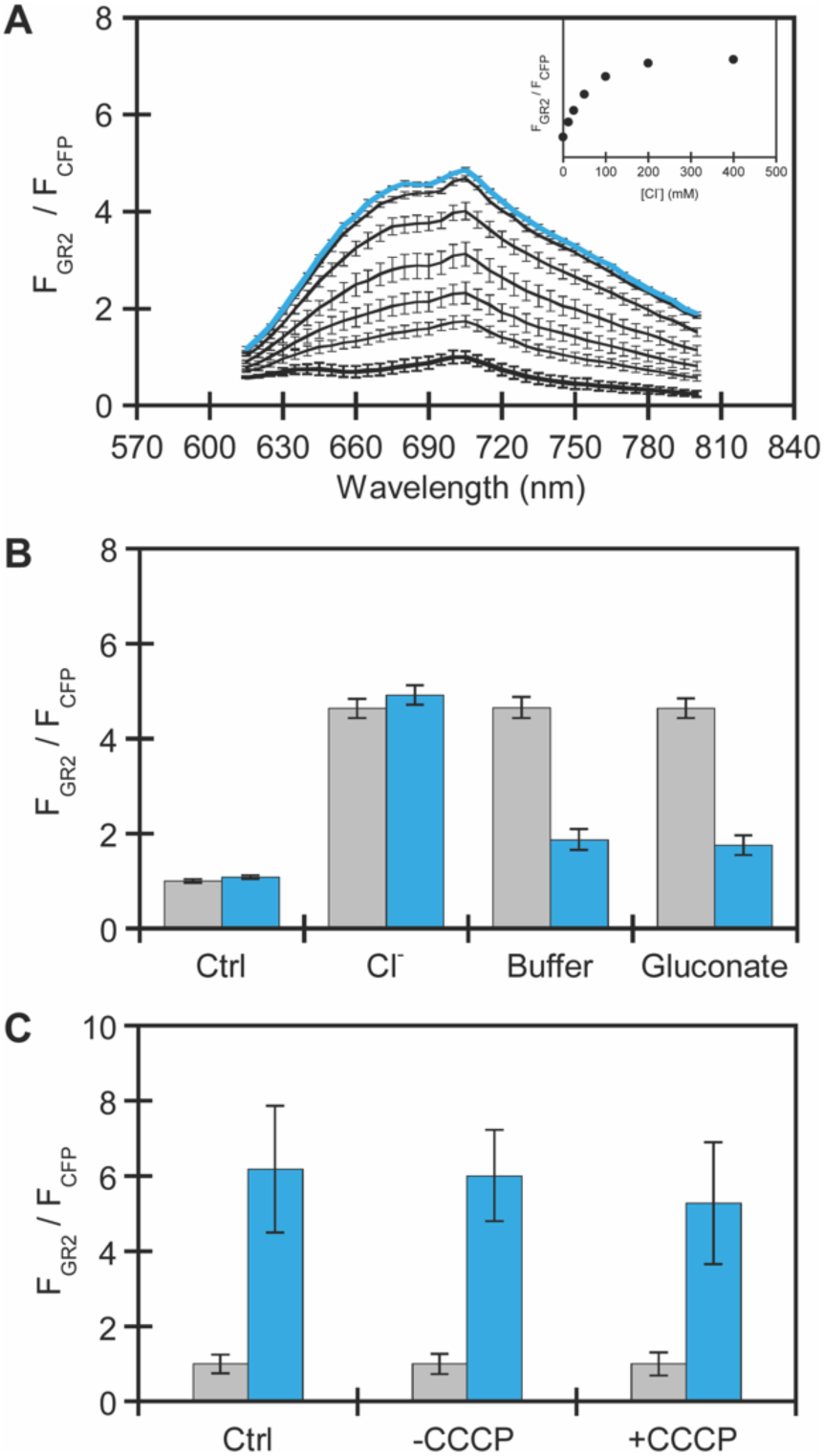
Characterization of GR2-CFP expressed in *E. coli* using bulk fluorescence plate reader experiments. (A) Normalized emission spectra of GR2-CFP in *E. coli* treated with 0 (black and bold), 12.5, 25, 50, 100, 200, and 400 mM (blue and bold) sodium chloride. The normalized emission response (F_GR2_/F_CFP_) is shown in the inset. (B) Normalized integrated emission response of GR2-CFP in *E. coli* first treated with buffer (Ctrl) or buffer with 400 mM sodium chloride (gray bars), followed by washing with buffer for the control sample and buffer with 400 mM sodium chloride, buffer only, or buffer with 400 mM sodium gluconate for the sodium chloride treated samples (blue bars). Spectra are shown in Figure S16–S18. (C) Normalized integrated emission response of GR2-CFP in *E. coli* treated with buffer, DMSO, or 30 μM CCCP in the presence of 0 mM or 400 mM sodium chloride. Spectra are shown in Figure S19–S21. All experiments were carried out in 50 mM sodium acetate buffer at pH 5. For the rhodopsin, the excitation was provided at 570 nm, and the emission was collected and integrated from 615–800 nm (F_GR2_). For CFP, the excitation was provided at 390 nm, and the emission was collected from 425–560 nm (F_CFP_). The GR2 emission spectra were normalized at each wavelength by the CFP emission intensity at 485 nm (F_CFP_). For each experiment, the average of nine biological replicates from three technical replicates with standard error of the mean is shown

Given the larger dynamic range, we further characterized GR2-CFP at pH 5. The chloride-dependent response can be reversed by washing cells with sodium acetate buffer (63% turn-off) or an equimolar amount sodium gluconate (65% turn-off) (Figure 4B, Figure S16–S18). This data suggests that GR2-CFP can dynamically sense changes in the extracellular and periplasmic chloride pool (Figure 4). Moreover, uncoupling of cell membrane potential with the protonophore carbonyl cyanide 3-chlorophenylhydrazone (CCCP) does not attenuate the fluorescence response (ΔF_GR_/F_CFP_ = 5.3 ± 1.6) in comparison to cells treated with a DMSO vehicle control (ΔF_GR_/F_CFP_ = 6.0 ± 1.2) (Figure 4C, Figure S19–S21). Thus, confirming that GR2-CFP does not sense membrane potential. Encouraged by these data, we used confocal fluorescence microscopy to validate GR2-CFP at the single cell level. Immobilization of *E. coli* on agarose pads at pH 5 shows that GR2 is non-fluorescent even though CFP is expressed in the cytosol (Figure 5, Figure S22, Figure S23). Upon supplementation with 400 mM sodium chloride, the GR2 signal is localized to the plasma membrane, and the fluorescence ratio increases by *ca*. 5.2-fold.

**Figure 5.**
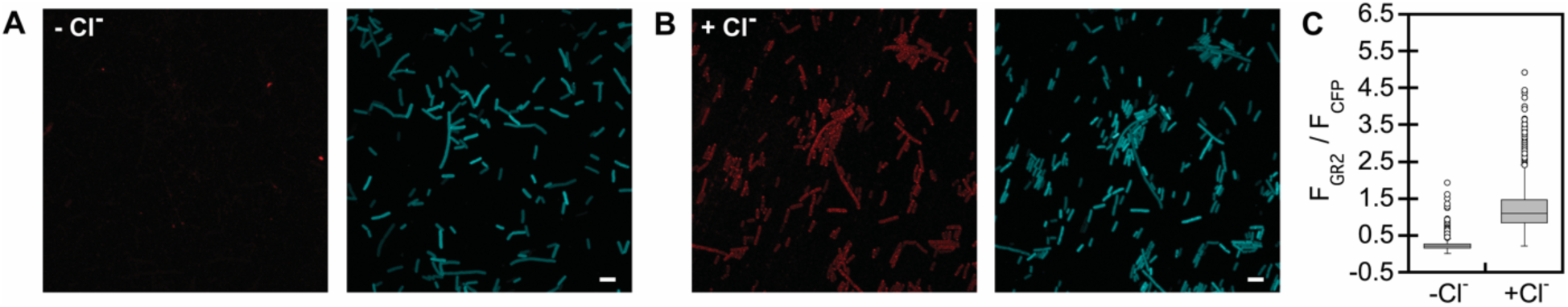
Confocal fluorescence microscopy experiments demonstrate that GR2-CFP is a ratiometric, turn-on fluorescent sensor for chloride at the single cell level in exogenously supplemented *E. coli*. Representative images are shown for *E. coli* expressing GR2-CFP on agarose pads with (A) 0 mM and (B) 400 mM sodium chloride in 50 mM acetate buffer at pH 5. For each panel, the GR2 emission (red) is on the left and CFP emission (cyan) is on the right (scale bar = 5 μm). (C) Boxplots show the normalized emission response (F_GR2_/F_CFP_) of each cell analyzed from three biological replicates (n = 2,583 regions of interest (ROIs) for 0 mM sodium chloride; n = 2,367 ROIs for 400 mM sodium chloride, Figure S22, Figure S23). The gray boxes correspond to the lower and upper quartile data with the minimum and maximum values extending below and above the box. The median values are indicated by the black lines in the gray boxes, and outliers are shown as open circles.

## Discussion

In this work, we show how the integration of evolutionary signals learned from extant sequences of the rhodopsin family can be used to guide the evolution of the engineered rhodopsin chloride sensor GR1-CFP into the improved variant GR2-CFP. Our integrated approach of computation and experimentation allows a significant reduction of mutational search space along the five proton-pumping pathway residues to accelerate the discovery process (Figure 1). To put this in perspective, a blind search of the possible variants would yield 19^5^ (≈ 2×10^6^) combinations to be screened. Our modeling approach first suggested residue positions with the largest score changes to be mutated in GR1-CFP. Using this, experimental site-saturation mutagenesis coupled with a high-throughput *in vivo* screening and selection enabled the identification of the amino acid substitutions at each position. The whole progress required the analysis of less than 95 (19×5) theoretical variants and 480 (96×5) experimental variants, which is significantly lower than sampling the complete space of variants (19^5^) that would be needed to engineer GR2-CFP.

Using bulk and single cell fluorescence assays, we show that GR2-CFP is a reversible, turn-on ratiometric fluorescent sensor for chloride. In comparison to GR1-CFP, GR2-CFP has a little to no background fluorescence from the rhodopsin (Figure 3, Figure 4A).^21^ This in combination with the larger dynamic range, translates into a *ca*. 3-fold higher ratio change for GR2-CFP versus GR1-CFP.^21^ However, the *K*_d_ for chloride binding to GR2-CFP (53 ± 14 mM) is comparable to that of GR1-CFP (42 ± 1 mM, Figure S10).^21^ Interestingly, even though we did not select for an expanded operational pH, the mutations along the proton-pumping pathway did result in measurable chloride sensing at pH 6 (Figure S14).

With respect to *wt*GR-CFP, GR2-CFP has five mutations: D121V, E132K, A84K, T125C, and V245I. In its native function, the D121 counterion position accepts a proton from the E132 donor and hydrogen bonds with T125, and residues A84, V245, and G261 line the proton-pumping pathway.^25^ As we previously reported, the non-natural D121V mutation in GR1-CFP eliminates proton-pumping activity and confers the fluorescence response to chloride at pH 5.^21,23^ In GR2-CFP, the enrichment of positively charged lysine residues at positions 132 and 84 provides a *ca*. 2-fold higher fluorescence ratio change relative to GR1-CFP (Figure 3, Figure S7, Figure S8). Mutation from a polar threonine to a larger, non-polar cysteine at position 125 in combination with the isoelectronic mutation from a valine to a larger isoleucine at position 245 increases the dynamic range from GR1-E132/A84K-CFP by an additional *ca*. 1.9-fold (Figure 3, Figure S7, Figure S8). It is interesting to note that the A84K and T125C mutations influence the non-natural chloride sensing function to a larger extent, but the order of mutagenesis was key to identify these improvements. This is perhaps unsurprising since both residues are adjacent to the putative chloride binding pocket and SBC (Figure 2A).^24^ Of the four mutations introduced into GR1-CFP, A84K is a non-natural substitution that has not been sampled in the natural evolution of the rhodopsin family (Figure S24).

Here, we only focused on a subset of proton-pumping pathway residues as an initial approach. However, examination of the top DI pairs reveals highly coupled residue pairs that merit further exploration to expand biosensor properties, including the turn-on fluorescence response, operational pH, and tuning of the excitation/emission wavelengths (Figure 2B). Thus, we are not necessarily limited to the residues along the proton-pumping pathway, and in principle, additional relevant sites could be elucidated without prior experimental knowledge. Efforts to expand our search space to new residues in GR2-CFP and other rhodopsins are currently in progress. Beyond the rhodopsin family, we will extend our collaborative approach to other chromophore-containing fluorescent proteins to identify, evolve, and design chloride-sensitive fluorescent proteins with new properties. With the knowledge gained, we will not only shape our understanding of the sequence level determinants that govern anion recognition in biology but will also generate an expanded set of sensors for cellular applications. To close, the use of evolutionary signals to engineer proteins with new properties has broad utility as we can readily extract direct knowledge from evolutionary processes that have taken long periods of time to fine tune the desired properties we see in Nature. We envision that learning and utilizing these signals as a guiding principle presents an exciting opportunity to intersect with existing and emerging protein engineering methods for fluorescent sensors and beyond.

## Methods

### Pre-processing of the rhodopsin family multiple sequence alignment (MSA)

The amino acid sequence of the wild-type proton pumping rhodopsin from *Gloeobacter violaceus* (*wt*GR) (UniProt ID: Q7NP59, residues 45–270) was used to search for homologous proteins in the rhodopsin family in UniProt Knowledgebase using the HMMER webserver v2.41.1.^55,56^ The threshold of expectation value (E) was set to 0.01 to filter and obtain 6,906 protein sequences that are like *wt*GR. Next, we excluded protein sequences containing consecutive gaps longer than 20% of the protein length (> 45 consecutive gaps) to reduce the noise from incomplete coverage. The amino acid composition at each proton-pumping pathway position in the sequence family was plotted (Figure S24). Then, re-weighted sequences with greater than 80% identity were filtered as previously described, leaving 879 effective sequences.^38^

### Direct coupling analysis of rhodopsin protein family

Direct coupling analysis (DCA) was performed to the pre-processed rhodopsin protein family sequences to infer a joint probability distribution that satisfies the statistical observations. Two types of parameters are estimated for this distribution: pairwise couplings *e*_*ij*_(*x*_*i*_, *x*_*j*_) and local biases *h*_*i*_(*x*_*i*_), where *i, j* indices refer to positions along the sequence and *x* refers to the amino acid at a specific position. Direct information (DI) values were computed from this distribution to quantify how the two sites are directly coupled.^41,57^ The top 300 high ranked DI pairs were plotted (Figure S2, Supplemental File 1). DI pairs related to proton-pumping involved residues were selected and plotted as a connectivity map (Figure 2B). For a given sequence L, the Hamiltonian score is defined as a sum of all the pairwise couplings *e*_*ij*_(*x*_*i*_, *x*_*j*_) and local biases *h*_*i*_(*x*_*i*_)^50^:

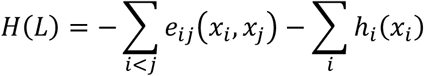

The fitness (functional change with respect to the parent) of mutants *σ*_*a*_ was predicted by the relative change of Hamiltonian score compared to the wild-type sequence (*wt*) as Δ*H*(*σ*_*a*_):

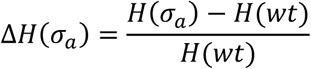

### Relative changes of the average Hamiltonian

To guide the iterative site-saturation mutagenesis, relative changes of the average Hamiltonian were calculated to evaluate the effect of changes for each proton-pumping pathway position (*σ*) involved.

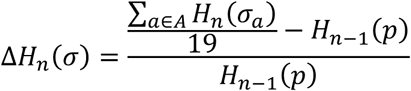

Where *n* is the number of mutation steps, *a* is the set of all the possible amino acid changes except the wild-type or parent amino acid. The variant with the most negative relative Hamiltonian score changes was selected as the parent (*H*_*n*−1_(*p*)) for the next step. We used *wt*GR as the parent in the first step for the single mutations.

### Homology model of GR1

The homology model of GR1 used in Figure 2 and Figure S2 was generated based on the crystal structure of *wt*GR (PDB ID: 6NWD). The homology model was generated using the MODELLER software.^58^

### General

All supplies and chemicals were purchased from Bio-Rad Laboratories, Sigma-Aldrich, Thermo Fisher Scientific, or VWR unless otherwise stated.

### Construction, expression, and screening of site-saturation mutagenesis (SSM) libraries

The pET-21a(+) based plasmid encoding GR1-CFP was generated in a prior study.^21^ Three forward primers and one reverse primer were synthesized using the 22c-trick (Sigma-Aldrich) to generate each SSM library as previously described (Table S2).^21,59^ *E. cloni* EXPRESS BL21 (DE3) cells (Lucigen) were transformed with each library and expressed in a 96-well deep well format.^21^ General methods for liquid handling and fluorescence plate reader screening at pH 5, 6, and 7 were adapted from methods to generate GR1-CFP.^21^ Variants with an increased turn-on fluorescence response with respect to the parent were re-screened at least in duplicate.^21^ If necessary, excitation and emission settings were altered for each experiment for optimal signal to noise. For the re-screen, variants were first selected based on improvement at pH 5. If improved variants were identified at pH 5, then the turn-on fluorescence response at pH 6 or 7 was used for selection. Based on this analysis, the plasmid DNA was isolated and sequenced as previously described.^21^

### Validation of improved variants

To compare the variants from each library, *E. cloni* EXPRESS BL21 (DE3) cells were transformed via electroporation and plated onto LB-Miller agar plates containing 100 μg/mL ampicillin (AMP). Three different colonies of each variant were tested for each of the three technical replicates. Single colonies were picked into 5 mL of LB media (10 g/L of tryptone and 5 g/L of yeast extract) containing 100 μg/mL AMP in a 14-mL culture tube and incubated overnight at 37 °C with shaking at 250 rpm. The next day, 150 μL of each overnight culture was diluted into 3 mL of fresh LB-AMP media and incubated for 2 h and 15 min at 30 °C with shaking at 230 rpm, followed by addition of 15 μL of 100 mM isopropyl β-D-1-thiogalactopyranoside (IPTG) in water and 15 μL of 2 mM all-*trans*-retinal (ATR) in ethanol for a final concentration of 500 μM and 10 μM, respectively. After 4 h, the cells were collected by centrifugation at 2,500*g* (5810 R, Eppendorf) for 5 min at room temperature. The resulting cell pellet was washed, resuspended, centrifuged, and resuspended in 500 μL of 50 mM sodium acetate buffer at pH 5. A 25 μL aliquot of the cell suspension was transferred to a well 96-well microtiter plate containing 175 μL 50 mM sodium acetate buffer at pH 5 or 175 μL 50 mM sodium acetate buffer at pH 5 with 457 mM sodium chloride. All spectra were collected on a Tecan plate reader. For the rhodopsins, the excitation was set to 570 nm (20 nm bandwidth), and the emission was collected from 630–760 nm (10 nm bandwidth, 10 nm step size, 100 gain, 25 flashes). Based on these spectra, the emission for the excitation scan was set to 710 nm (20 nm bandwidth), and the excitation was collected from 490–630 nm (20 nm bandwidth, 100 gain, 10 nm step size, 25 flashes). For CFP, the excitation was set to 390 nm (10 nm bandwidth), and the emission was collected from 435–520 nm (20 nm bandwidth, 5 nm step size, 55 gain, 10 flashes). For all rhodopsin emission spectra, the emission intensity at each wavelength was normalized to the emission intensity of CFP at 485 nm. The normalized emission spectra were then averaged across all twelve biological replicates. These spectra are reported with the standard error of the mean (Figure S7, Figure S8). To determine the turn-on response, the emission spectra of GR2 and CFP were first integrated in Kaleidagraph v4.5 (Synergy Software) from 630–760 nm and 435–520 nm, respectively. The integrated emission intensity from the rhodopsin (F_GR_) was then normalized to the integrated emission intensity from CFP (F_CFP_). The turn-on emission responses were calculated by dividing the F_GR_/F_CFP_ ratios for 400 mM sodium chloride by 0 mM sodium chloride. The average of all twelve biological replicates across three technical replicates with standard error of the mean is reported (Figure 3).

### SDS-PAGE and Western blot analysis of GR2-CFP

*E. cloni* EXPRESS BL21 (DE3) cells were transformed with the plasmid DNA encoding for the GR2-CFP construct via electroporation and plated onto LB-Miller agar plates containing 100 μg/mL AMP. Single colonies were picked into 5 mL of LB media containing 100 μg/mL AMP in a 14-mL culture tube and incubated overnight at 37 °C with shaking at 250 rpm. The following morning, the overnight culture was diluted 1:20 into 25 mL of LB media containing 100 μg/mL AMP in 125-mL baffled flasks. The cultures were then incubated for 2 h at 30 °C with shaking at 230 rpm. Following the 2 h incubation period, protein expression was induced with 122 μL of 100 mM IPTG and 23 μL of 12 mM ATR for a final concentration of 500 μM and 10 μM, respectively. Non-induced cultures served as negative controls. The cultures were incubated for 4 h at 30 °C with shaking at 230 rpm. After the final incubation period, a 6 mL portion of each cell culture was collected by centrifugation at 2,500*g* (5810 R, Eppendorf) for 5 min at 4 °C. Following this, the cells were resuspended in 500 μL of lysis buffer containing 20 mM Tris buffer at pH 7.5 with 200 mM sodium chloride, 5 mM magnesium chloride, 30 μg/mL deoxyribonuclease I, and two cOmplete protease inhibitor tablets. The resuspended cell pellets were lysed with a pellet mixer equipped with a cordless motor in 30 s on-off intervals for 3 min on ice. Following homogenization, the lysates were clarified via centrifugation at 5,000*g* (5424 R, Eppendorf) for 3 min at 4 °C.

The total protein concentration of each sample was determined using a commercial bicinchoninic acid protein assay. For each sample, 18 μL of 0.5 mg/mL protein was combined with 6 μL of 4X Laemmli sample buffer) supplemented with 10% beta-mercaptoethanol and 50 mM Tris buffer at pH 7.5 for a total volume of 24 μL. The protein-buffer mixtures were heated at 37 °C for 10 min, and then 15 μL of each sample were loaded onto a 12% Mini-PROTEAN TGX acrylamide gel alongside 5 μL of a prestained protein ladder.

The gel was run in 1X Tris-Glycine-SDS running buffer containing 25 mM Tris at pH 8.3 with 192 mM glycine and 0.1% SDS at 150 V for 1 h. The separated proteins were then transferred to a methanol-activated polyvinylidene difluoride (PVDF) membrane utilizing the Bio-Rad Trans-Blot Turbo Transfer system and kit with the seven-minute mixed molecular weight transfer setting. All incubation and wash steps were performed with orbital shaking. Following the protein transfer, the membrane was blocked with 5% milk in 1X Tris-Buffered Saline containing 0.1% Tween20 (TBST) for 1 h at room temperature. After blocking, the membrane was incubated with 3 μL of 1 mg/mL 6x-His tag monoclonal antibody in 5% milk in TBST at 4 °C. The next day, the primary antibody was removed, and the membrane was washed with TBST (3 × 5 min) and then incubated with 15 μL of 2 mg/mL goat anti-mouse horse radish peroxidase conjugated secondary antibody in 5% milk in TBST at 4 °C for 2 h. The secondary antibody solution was decanted, and then the membrane was washed with TBST (3 × 5 min). For visualization, the membrane was incubated in 2 mL of the Immobilon Classico Western HRP substrate for 1 min and imaged on a Bio-Rad ChemiDoc imaging station utilizing the colorimetric setting for the ladder and the chemiluminescent setting for protein imaging. The resulting images were overlayed and analyzed with the Bio-Rad ImageLab software (Figure S9).

### Characterization of GR2-CFP in *E. coli* using bulk fluorescence measurements

*E. cloni* EXPRESS BL21 (DE3) cells were freshly transformed with plasmid DNA encoding for the GR2-CFP construct via electroporation and plated onto LB-Miller agar plates containing 100 μg/mL AMP. For each biological replicate, a single colony was picked into 5 mL of LB media with 100 μg/mL AMP in a 14-mL culture tube and grown overnight at 37 °C with shaking at 250 rpm. The following day, the overnight culture was diluted 1:20 in 25 mL of LB media containing 100 μg/mL AMP in a 125-mL baffled flask and incubated for 2 h at 30 °C with shaking at 230 rpm. Following this, protein expression was induced with 122 μL of 100 mM IPTG and 23 μL of 12 mM ATR in ethanol for a final concentration of 500 μM and 10 μM, respectively. The induced culture was incubated for 4 h at 30 °C with shaking at 230 rpm, and then 6 mL of the culture was collected by centrifugation at 2,500*g* (5810 R, Eppendorf) for 5 min at room temperature.

Chloride titrations with GR2-CFP expressing cells were carried out to determine the dynamic ranges and apparent dissociation constants (*K*_d_) for chloride. The resulting cell pellet was resuspended in 500 μL of 50 mM sodium acetate buffer at pH 5 or 50 mM sodium phosphate buffer at pH 6 and pH 7, and then re-collected by centrifugation and resuspended in 500 μL of the corresponding buffer. In a 96-well microtiter plate, a 25 μL portion of the cell suspension was further diluted with 175 μL of 50 mM sodium acetate buffer at pH 5 or sodium phosphate buffer at pH 6 and pH 7 and the corresponding buffer containing 14.3, 28.5, 57, 114.3, 228.5, 457, or 685.7 sodium chloride to the final concentrations of 0, 12.5, 25, 50, 100, 200, 400, and 600 mM sodium chloride. All spectra were collected on a Tecan plate reader. For the rhodopsin, the excitation was set to 570 nm (10 nm bandwidth), and the emission was collected from 615–800 nm (10 nm bandwidth, 5 nm step size, 150 gain, 25 flashes). For CFP, excitation was set to 390 nm (10 nm bandwidth), and emission was collected from 425–560 nm (20 nm bandwidth, 5 nm step size, 100 gain, 25 flashes). For all rhodopsin spectra, the emission intensity at each wavelength was normalized to the emission intensity of CFP at 485 nm. The normalized emission spectra were then averaged across all nine biological replicates. These spectra are reported with the standard error of the mean (Figure 4A, Figure S10–S15). To determine the turn-on responses and *K*_d_s, the emission spectra of GR2 and CFP were first integrated in Kaleidagraph v4.5 (Synergy Software) from 615–800 nm and 425–560 nm, respectively. The integrated emission intensity from the rhodopsin (F_GR2_) was then normalized to the integrated emission intensity from CFP (F_CFP_). The turn-on emission responses were calculated by dividing the F_GR2_/F_CFP_ ratios for 400 mM sodium chloride by 0 mM sodium chloride. To determine the *K*_d_s, the emission response of GR2-CFP was calculated with the following equation where F_obs_ is the observed F_GR2_/F_CFP_ at a given concentration of sodium chloride, and F_min_ and F_max_ correspond to F_GR2_/F_CFP_ at 0 mM and 600 mM sodium chloride, respectively: F=(F_obs_-F_min_)/(F_max_-F_min_). These emission responses were plotted versus the concentration of sodium chloride and fitted to the following equation to determine the *K*_d_ where F is observed F_GR2_/F_CFP_ at a given concentration of sodium chloride and B_max_ represents the fluorescence emission at saturation: F=([Cl^-^] × B_max_)/(*K*_d_ + [Cl^-^]). The average of all nine biological replicates across three technical replicates with standard error of the mean is reported (Figure S10, Figure S14, Figure S15).

Control experiments were also performed to observe any background fluorescence of the *E. coli* cells that may affect the GR2-CFP emission signal with the following expression conditions. One culture tube was expressed with the same IPTG and ATR conditions as described above. The second culture tube was treated with IPTG in the absence of ATR, the third culture tube was treated only with ATR and no IPTG, and the fourth culture tube did not receive either IPTG or ATR. Each of the cultures was then incubated at 30 °C with shaking at 230 rpm for 4 h. Following protein expression as described above, the cells were then harvested and scanned on a Tecan plate reader using the same conditions as described above. For all rhodopsin spectra, the emission intensity at each wavelength was normalized to the emission intensity of CFP at 485 nm. The normalized spectra of all nine biological replicates across three technical replicates with standard error of the mean is reported (Figure S11–S13).

To test the reversibility of GR2-CFP, cultures of *E. cloni* expressing GR2-CFP were prepared as described above. The resulting cell pellet was resuspended in 400 μL of 50 mM sodium acetate buffer at pH 5. In 1.5-mL centrifuge tubes, a 75 μL portion of the cell suspension was further diluted with 525 μL of 50 mM sodium acetate buffer at pH 5 (control) or buffer containing 457 mM sodium chloride to a final concentration of 0 mM or 400 mM sodium chloride. After mixing and incubating at room temperature for 10 min, a 200 μL portion of each cell suspension was transferred to a 96-well microtiter plate, and spectra were collected as described above. The remaining cell suspensions were collected by centrifugation at 500*g* (5424 R, Eppendorf) for 10 min at room temperature and resuspended as follows: the control cell pellet was resuspended in 400 μL of 50 mM sodium acetate buffer at pH 5, and the remaining cell pellets were resuspended in 400 μL of buffer or buffer containing 400 mM sodium chloride or 400 mM sodium gluconate. After mixing and incubating at room temperature for 10 min, a 200 μL portion of each cell suspension was transferred to a 96-well microtiter plate, and spectra were collected and analyzed as described above. The average of all nine biological replicates across three technical replicates with standard error of the mean is reported (Figure 4B, Figure S16–S18).

To validate that the turn-on response of GR2-CFP was not dependent on membrane potential, GR2-CFP expressing cells were treated with CCCP. Cultures of *E. cloni* expressing GR2-CFP were prepared as described above. The cells were collected by centrifugation at 2,500*g* (5810 R, Eppendorf) for 5 min at room temperature, and the resulting cell pellet was resuspended in 80 μL of 50 mM sodium acetate buffer at pH 5. In a 96-well microtiter plate, a 10 μL portion of the cell suspension was further diluted with 175 μL of 50 mM sodium acetate buffer at pH 5 or buffer containing 457 mM sodium chloride to a final concentration of 0 mM or 400 mM sodium chloride. Each well was treated with 15 μL of 50 mM sodium acetate buffer at pH 5, 4% DMSO (v/v), or 400 μM CCCP in DMSO in 50 mM sodium acetate buffer at pH 5 to a final concentration of 0.3% DMSO (v/v) or 30 μM CCCP. The microtiter plate was incubated in the plate reader at 30 °C for 30 min with shaking for 30 s every 5 min, and the spectra were collected and analyzed as described above. The average of all nine biological replicates across three technical replicates with standard error of the mean is reported (Figure 4C, Figure S19–S21).

### Validation of GR2-CFP at the single cell level in *E. coli* with fluorescence microscopy

Three-mL cultures of *E. cloni* expressing GR2-CFP were expressed as described above for the 25-mL cultures. The cells were harvested by centrifugation at 2,500*g* (5810 R, Eppendorf) for 5 min at room temperature. The resulting cell pellet was resuspended in 200 μL of 50 mM sodium acetate buffer at pH 5 and 25 μL of the resuspension was further diluted into 175 μL of 50 mM sodium acetate buffer at pH 5 or buffer containing 457 mM sodium chloride to a final concentration of 0 mM or 400 mM sodium chloride. The cells were gently mixed, and 3–6 μL were transferred to 1.5% agarose pads containing 0 mM or 400 mM sodium chloride and imaged on 35 mm dishes with a 20 mm glass-bottom coverslip (No. 1.5, MatTek) as previously described.^21^

The fluorescence images were acquired as Z-stacks using a confocal laser scanning microscope (FV3000RS, Olympus) with a 100X silicon immersion objective.^21^ Briefly, excitation for GR2 was provided at 561 nm, and the emission was collected from 630–730 nm. Excitation for CFP was provided at 405 nm, and emission was collected from 450–550 nm. For each Z-stack, the maximum intensity Z-projection for the CFP channel was first duplicated and then the default threshold settings were applied using the software Fiji Is Just ImageJ (Fiji) v2.0.^60^ Masks were created using the thresholded CFP image, and the Fiji Analyze Particles function was used to select regions of interest (ROIs) that were greater than 3 pixels with circularity between 0 and 1. Overlapping cells were considered as one ROI. For each sample, the ROIs were transferred to the unprocessed GR2 and CFP maximum intensity Z-projections, and the median fluorescence intensities were measured. ROIs that did not appear in both the GR2 and CFP channels were manually excluded. The GR2 median fluorescence intensity was normalized to the CFP median fluorescence intensity (Figure 5, Figure S22, Figure S23). At least five different fields were analyzed for the three biological replicates (n = 2,583 ROIs for 0 mM sodium chloride; n = 2,367 ROIs for 400 mM sodium chloride).

## Acknowledgements

We acknowledge Dr. Ved Prakash from the Olympus Discovery Center at UT Dallas for expert technical assistance with confocal microscopy experiments. F.M. acknowledges support from the National Science Foundation (MCB-1943442) and the National Institute of General Medical Sciences of the National Institutes of Health (R35GM133631). S.C.D. acknowledges support from The University of Texas at Dallas, the Welch Foundation (AT-1918-20170325, AT-2060-20210327), and the National Institute of General Medical Sciences of the National Institutes of Health (R35GM128923). This manuscript is the sole responsibility of the authors and does not represent the views of the funding sources.

