## Supporting Information for "Coupling a Live Cell Directed Evolution Assay with Coevolutionary Landscapes to Engineer an Improved Fluorescent Rhodopsin Chloride Sensor"

```
BR    ------------------------------QAQITGRPEWIWLALGT--ALMGLGTLYFL  28
GR    MLMTVFSSAPELALLGSTFAQVDPSNLSVSDSLTYGQFNLVYNAFSFAIAAMFASALFFF  60
                                    ::   *: : :: *:.   * *  .:*:*:

BR    VKGMGVSDPDAKKFYAITTLVPA----IAFTMYLSMLLGYGLTMVPFGG-----EQNPIY  79
GR    SAQALV-----GQRYRLALLVSAIVVSIAGYHYFRIFNSWDAAYVLENGVYSLTSEKFND  115
           *      : * :: ** *    **   *: :: .:. : *  .*     .::
                                  84

BR    WARYADWLFTTPLLLLDLALLVD----ADQGTILALVGADGIMIGTGLVGALTKVYSYRF  135
GR    AYRYVDWLLTVPLLLVETVAVLTLPAKEARPLLIKLTVASVLMIATGYPGEISDDITTRI  175
        **.***:*.****:: . ::       :  :: *. *. :**.**  * ::.  : *:
          121 125     132

BR    VWWAISTAAMLYILYVLFFGFTSKAESMRPEVASTFKVLRNVTVVLWSAYPVVWLIGSEG  195
GR    IWGTVSTIPFAYILYVLWVELSRSLVRQPAAVQTLVRNMRWLLLLSWGVYPIAYLLPMLG  235
      :* ::**  : ******:. :: .       * : .: :* : :: *..**:.:*:   *

BR    AGIVPLNI-ETLLFMVLDVSAKVGFGLILLRSRAIF--GEAEAPEPSAGDGAAATS----  248
GR    VSGTSAAVGVQVGYTIADVLAKPVFGLLVFAIALVKTKADQESSEPHAAIGAAANKSGGS  295
      .. .    :    : : : ** **  ***:::    :    .: *: ** *. ****..
              245              261

BR    ---                                                           248
GR    LIS                                                           298
```

**Figure S1.** Multiple sequence alignment of bacteriorhodopsin (BR, PDB ID: 1FBB) and *Gloeobacter violaceus* rhodopsin (*wt*GR, PDB ID: 6NWD) generated using Clustal Omega.[1] Sequences were derived from the respective Protein Data Bank files. Residues along the proton-pumping pathway selected for this study are shown in red and labeled with the position number from *wt*GR.

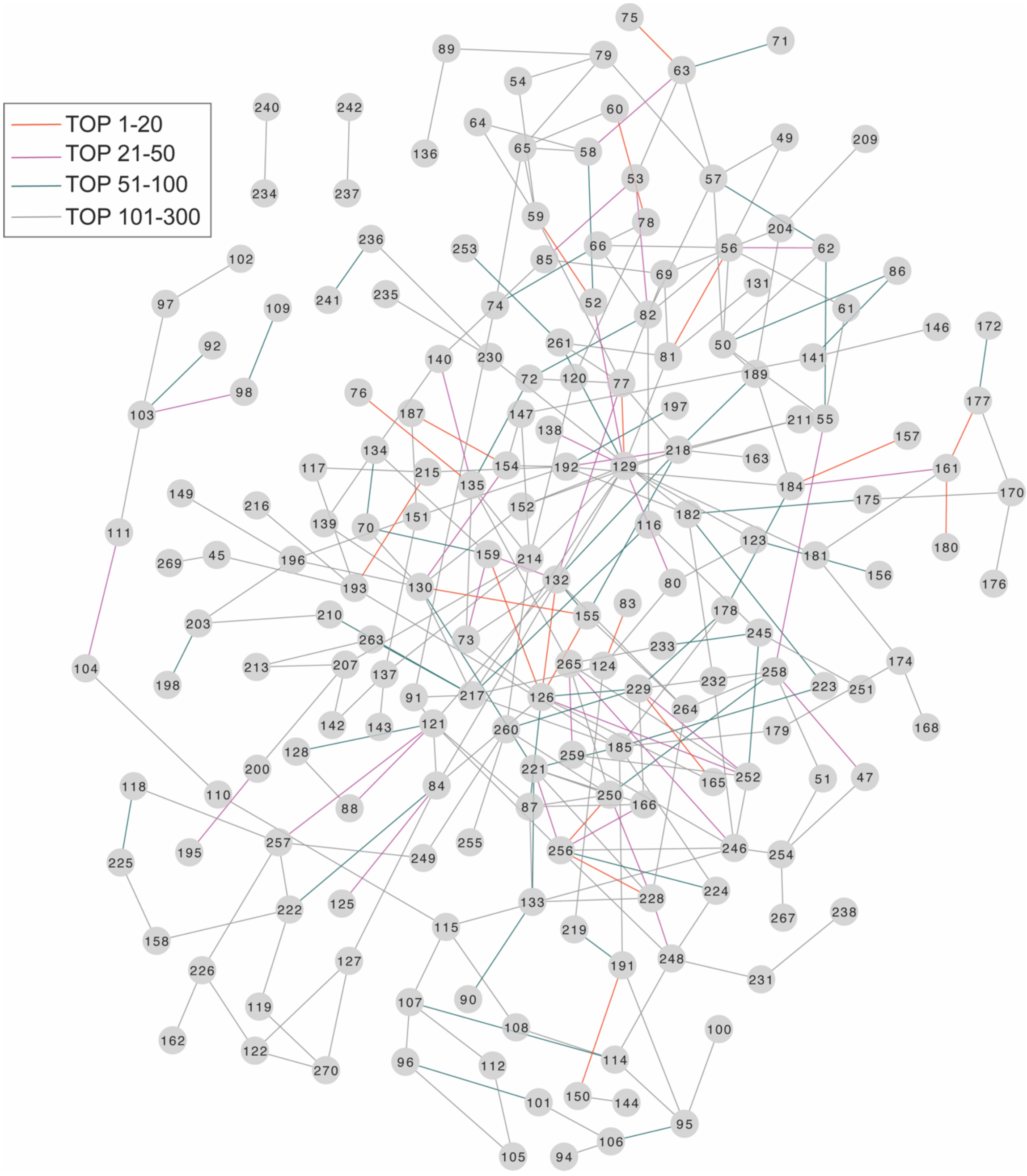


**Figure S2.** Connectivity map for the top 300 direct information (DI) pairs. Residues that maintain strong direct evolutionary coupling strength are shown in the map with position numbers mapped to *wt*GR (PDB ID: 6NWD) in gray circles. Each DI pair is linked with a line. The colors of lines represent the rank of the strength of the couplings for the DI pairs.

**Supplemental File 1.** Table of the top 300 DI pairs.

**Table S1.** Distances for the top DI pairs for the selected proton-pumping pathway residues. The shortest distances between the sidechains are reported.

| Residue | Amino acid | Paired residue | Amino acid | Distance (Å) |
|---|---|---|---|---|
| 84 | ALA | 121 | VAL | 5.2 |
| | | 125 | THR | 4.6 |
| | | 126 | VAL | 8.9 |
| | | 127 | PRO | 11.3 |
| | | 129 | LEU | 8.4 |
| | | 222 | TRP | 12.9 |
| 121 | VAL | 88 | TYR | 5.6 |
| | | 84 | ALA | 5.2 |
| | | 91 | ILE | 4.8 |
| | | 87 | HIS | 2.9 |
| | | 128 | LEU | 7.5 |
| | | 132 | GLU | 11.3 |
| | | 256 | ALA | 10.1 |
| | | 257 | LYS | 5.7 |
| 125 | THR | 84 | ALA | 4.6 |
| 132 | GLU | 73 | ALA | 4.1 |
| | | 77 | SER | 2.8 |
| | | 121 | VAL | 11.3 |
| | | 126 | VAL | 6.9 |
| | | 155 | SER | 11.0 |
| | | 159 | ILE | 14.5 |
| | | 218 | LEU | 9.4 |
| | | 264 | VAL | 3.8 |
| 245 | VAL | 116 | ALA | 11.2 |
| | | 233 | MET | 8.8 |
| | | 251 | ILE | 7.0 |
| | | 252 | ALA | 7.8 |
| 261 | GLY | 77 | SER | 4.4 |
| | | 81 | VAL | 8.3 |
| | | 129 | LEU | 6.9 |
| | | 253 | ASP | 8.2 |

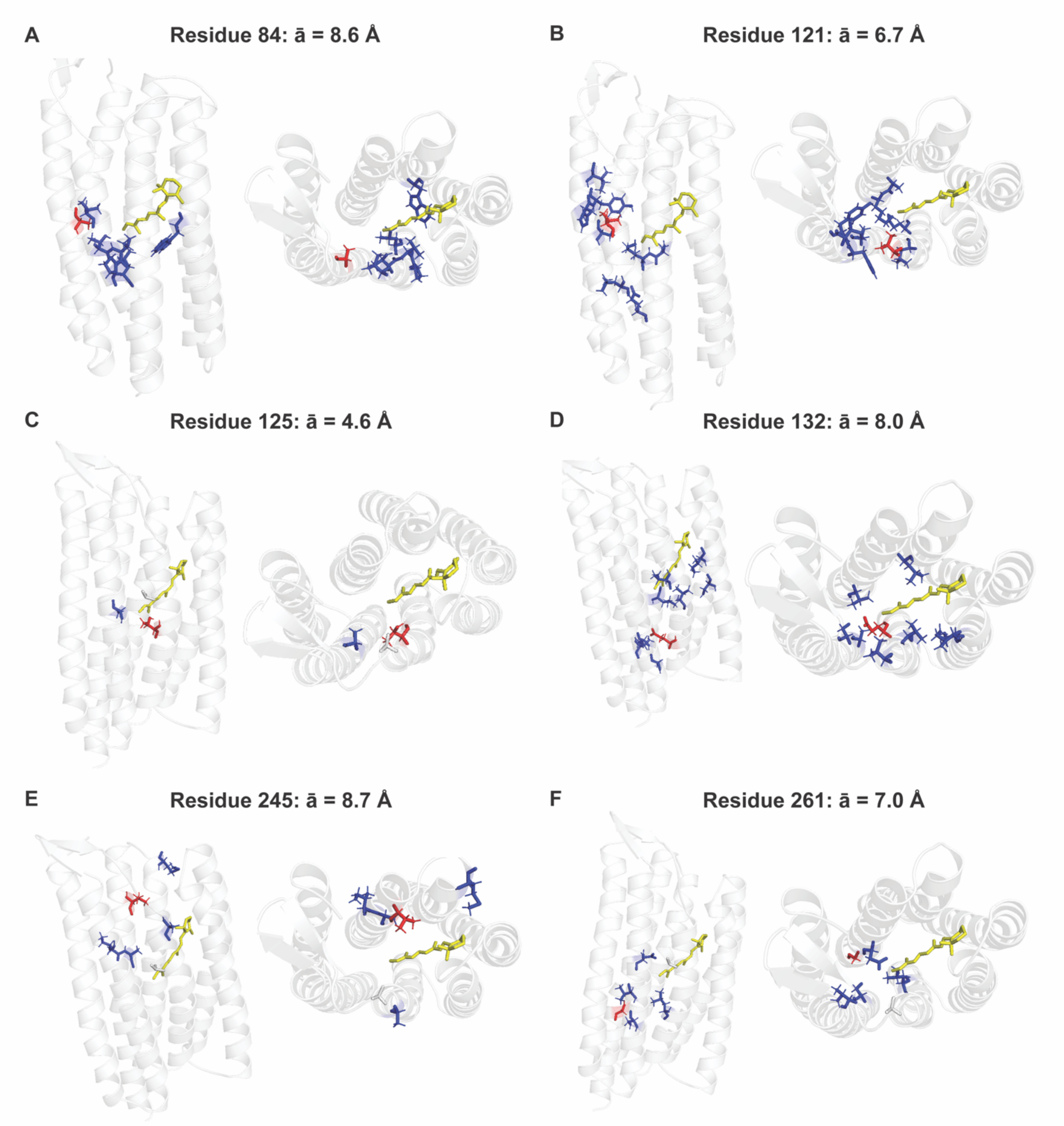


**Figure S3.** (A–F) Comparison of the residues along the proton-pumping pathway (red sticks) with the corresponding coevolved residues (blue sticks) for the top DCA pairs shown in Figure 2. In each panel, the side (left) and top (right) views are shown of the homology model for the rhodopsin chloride sensor GR1 only with the Schiff base chromophore (yellow sticks). The average distances ($\bar{a}$) from the proton-pumping pathway residues (red sticks) to the corresponding co-evolved residues (blue sticks) are listed in Table S1.

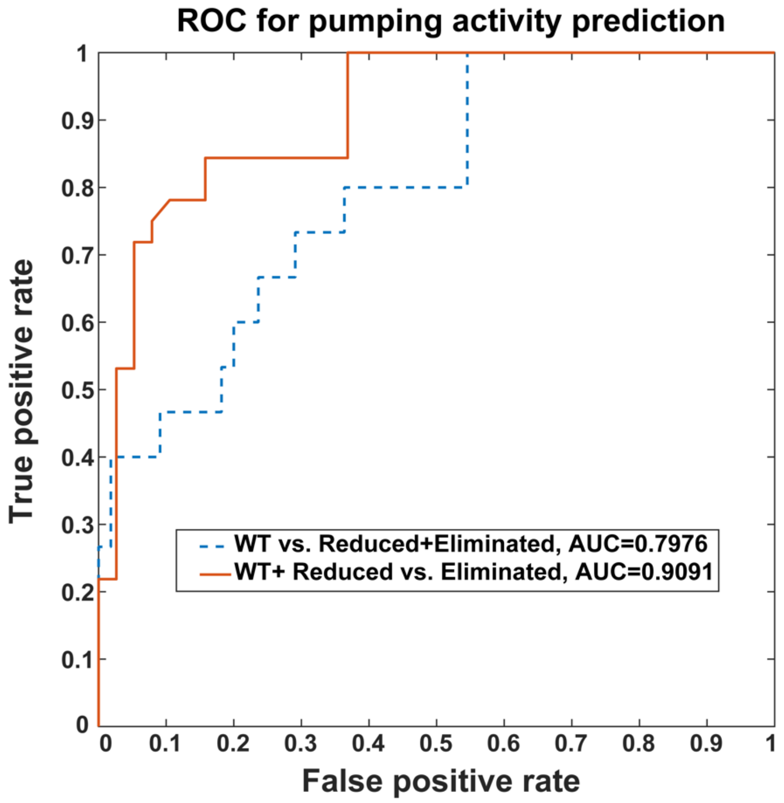


**Figure S4**. Receiver operating characteristic (ROC) curve for the Hamiltonian score prediction. Based on the proton-pumping activity, two definitions were used to define a positive. For the first definition, only variants with proton-pumping activity like *wt*GR were defined as positive. This curve is shown as a red solid line. The area under this ROC curve is 0.7976. For the second definition, only variants that lost proton-pumping activity were defined as negative. This curve is shown as a blue dashed line. The area under this ROC curve is 0.9091.

**Table S2.** List of the primers used to generate the site-saturation mutagenesis (SSM) library at each position. The mutation site is in red.

| Description | Primer (5' to 3') |
|---|---|
| 132 SSM Forward | GTGCCTCTGTTGCTGGTGNDTACAGTGGCAGTGCTGA |
| | GTGCCTCTGTTGCTGGTGVHGACAGTGGCAGTGCTGA |
| | GTGCCTCTGTTGCTGGTGTGGACAGTGGCAGTGCTGA |
| 132 SSM Reverse | CACCAGCAACAGAGGCACGGTCAACAGCCAAACC |
| 84 SSM Forward | GCAATTGTTGTGAGTATCNDTGGGTACCACTACTTTC |
| | GCAATTGTTGTGAGTATCVHGGGGTACCACTACTTTC |
| | GCAATTGTTGTGAGTATCTGGGGGTACCACTACTTTC |
| 84 SSM Reverse | GATACTCACAACAATTGCTGAAACAAGCAAGGCC |
| 125 SSM Forward | TATGTGGTTTGGCTGTTGNDTGTGCCTCTGTTGCTGG |
| | TATGTGGTTTGGCTGTTGVHGGTGCCTCTGTTGCTGG |
| | TATGTGGTTTGGCTGTTGTGGGTGCCTCTGTTGCTGG |
| 125 SSM Reverse | CAACAGCCAAACCACATAGCGGTAGGCGTCGTTG |
| 245 SSM Forward | ACGTCCGCGGCTGTCGGCNDTCAGGTTGGCTATACGA |
| | ACGTCCGCGGCTGTCGGCVHGCAGGTTGGCTATACGA |
| | ACGTCCGCGGCTGTCGGCTGGCAGGTTGGCTATACGA |
| 245 SSM Reverse | GCCGACAGCCGCGGACGTACCGGATACTCCAAGC |
| 261 SSM Forward | CTGGCGAAGCCTGTATTTNDTCTTCTAGTCTTCGCGA |
| | CTGGCGAAGCCTGTATTTVHGCTTCTAGTCTTCGCGA |
| | CTGGCGAAGCCTGTATTTTGGCTTCTAGTCTTCGCGA |
| 261 SSM Reverse | AAATACAGGCTTCGCCAGCACGTCTGCGATCGTA |

CATATGTTGATGACCGTATTTTCTTCTGCACCTGAACTTGCCCTTCTCGGATCAACCTTTGC
CCAGGTCGATCCTTCAAACTTATCGGTCTCAGATTCGCTGACCTATGGTCAGTTCAATCTG
GTTTACAACGCTTTCTCGTTTGCCATCGCGGCAATGTTCGCATCTGCCCTCTTCTTCTTCAG
CGCTCAGGCACTCGTCGGTCAACGATACCGGTTGGCCTTGCTTGTTTCAGCAATTGTTGTG
AGTATCAAGGGGTACCACTACTTTCGGATCTTCAATAGTTGGGATGCTGCCTACGTTCTGG
AGAATGGCGTGTATTCCCTGACTAGCGAAAAATTCAACGACGCCTACCGCTATGTGGTTTG
GCTGTTGTGTGTGCCTCTGTTGCTGGTGAAGACAGTGGCAGTGCTGACGTTGCCTGCAAA
GGAGGCAAGACCCTTGCTGATCAAACTGACGGTGGCTTCAGTTCTGATGATTGCCACGGG
CTACCCCGGCGAGATTTCTGACGACATTACGACTCGCATCATCTGGGGTACGGTCAGCAC
GATTCCCTTCGCCTACATCCTCTATGTGTTGTGGGTCGAACTGTCCAGGTCCCTTGTCCGC
CAGCCCGCTGCTGTACAAACCCTGGTCCGCAACATGCGGTGGCTGCTGTTGCTCTCCTGG
GGTGTTTACCCGATCGCATACCTTCTACCCATGCTTGGAGTATCCGGTACGTCCGCGGCT
GTCGGCATTCAGGTTGGCTATACGATCGCAGACGTGCTGGCGAAGCCTGTATTTGGTCTT
CTAGTCTTCGCGATTGCACTCGTGAAAACAAAAGCAGATCAAGAAAGCAGTGAACCACATG
CCGCAATAGGTGCTGCTGCAAATAAATCGGGAGGCAGTCTTATCTCCGCGGCCGCAATGG
TGAGCAAGGGCGAGGAGCTGTTCACCGGGGTGGTGCCCATCCTGGTCGAGCTGGACGGC
GACGTAAACGGCCACAAGTTCAGCGTGTCCGGCGAGGGCGAGGGCGATGCCACCTACGG
CAAGCTGACCCTGAAGTTCATCTGCACCACCGGCAAGCTGCCCGTGCCCTGGCCCACCCT
CGTGACCACCCTGACCTGGGGCGTGCAGTGCTTCAGCCGCTACCCCGACCACATGAAGC
AGCACGACTTCTTCAAGTCCGCCATGCCCGAAGGCTACGTCCAGGAGCGCACCATCTTCT
TCAAGGACGACGGCAACTACAAGACCCGCGCCGAGGTGAAGTTCGAGGGCGACACCCTG
GTGAACCGCATCGAGCTGAAGGGCATCGACTTCAAGGAGGACGGCAACATCCTGGGGCA
CAAGCTGGAGTACAACTACATCAGCCACAACGTCTATATCACCGCCGACAAGCAGAAGAAC
GGCATCAAGGCCAACTTCAAGATCCGCCACAACATCGAGGACGGCAGCGTGCAGCTCGC
CGACCACTACCAGCAGAACACCCCCATCGGCGACGGCCCCGTGCTGCTGCCCGACAACC
ACTACCTGAGCACCCAGTCCGCCCTGAGCAAAGACCCCAACGAGAAGCGCGATCACATGG
TCCTGCTGGAGTTCGTGACCGCCGCCCTCGAGCACCACCACCACCACCACTGA

**Figure S5.** Nucleotide sequence encoding GR2-CFP in the pET-21a(+) vector. The following regions are highlighted: restriction sites and the *A* nucleotide (orange), GR2 (black), mutation sites in GR2 (red), CFP (blue), His-tag (purple), and stop codon (green).

```
Met Leu Met Thr Val Phe Ser Ser Ala Pro Glu Leu Ala Leu Leu Gly Ser Thr Phe Ala 20
Gln Val Asp Pro Ser Asn Leu Ser Val Ser Asp Ser Leu Thr Tyr Gly Gln Phe Asn Leu 40
Val Tyr Asn Ala Phe Ser Phe Ala Ile Ala Ala Met Phe Ala Ser Ala Leu Phe Phe Phe 60
Ser Ala Gln Ala Leu Val Gly Gln Arg Tyr Arg Leu Ala Leu Leu Val Ser Ala Ile Val 80
Val Ser Ile Lys Gly Tyr His Tyr Phe Arg Ile Phe Asn Ser Trp Asp Ala Ala Tyr Val 100
Leu Glu Asn Gly Val Tyr Ser Leu Thr Ser Glu Lys Phe Asn Asp Ala Tyr Arg Tyr Val 120
Val Trp Leu Leu Cys Val Pro Leu Leu Leu Val Lys Thr Val Ala Val Leu Thr Leu Pro 140
Ala Lys Glu Ala Arg Pro Leu Leu Ile Lys Leu Thr Val Ala Ser Val Leu Met Ile Ala 160
Thr Gly Tyr Pro Gly Glu Ile Ser Asp Asp Ile Thr Thr Arg Ile Ile Trp Gly Thr Val 180
Ser Thr Ile Pro Phe Ala Tyr Ile Leu Tyr Val Leu Trp Val Glu Leu Ser Arg Ser Leu 200
Val Arg Gln Pro Ala Ala Val Gln Thr Leu Val Arg Asn Met Arg Trp Leu Leu Leu Leu 220
Ser Trp Gly Val Tyr Pro Ile Ala Tyr Leu Leu Pro Met Leu Gly Val Ser Gly Thr Ser 240
Ala Ala Val Gly Ile Gln Val Gly Tyr Thr Ile Ala Asp Val Leu Ala Lys Pro Val Phe 260
Gly Leu Leu Val Phe Ala Ile Ala Leu Val Lys Thr Lys Ala Asp Gln Glu Ser Ser Glu 280
Pro His Ala Ala Ile Gly Ala Ala Ala Asn Lys Ser Gly Gly Ser Leu Ile Ser Ala Ala 300
Ala Met Val Ser Lys Gly Glu Glu Leu Phe Thr Gly Val Val Pro Ile Leu Val Glu Leu 320
Asp Gly Asp Val Asn Gly His Lys Phe Ser Val Ser Gly Glu Gly Glu Gly Asp Ala Thr 340
Tyr Gly Lys Leu Thr Leu Lys Phe Ile Cys Thr Thr Gly Lys Leu Pro Val Pro Trp Pro 360
Thr Leu Val Thr Thr Leu Thr Trp Gly Val Gln Cys Phe Ser Arg Tyr Pro Asp His Met 380
Lys Gln His Asp Phe Phe Lys Ser Ala Met Pro Glu Gly Tyr Val Gln Glu Arg Thr Ile 400
Phe Phe Lys Asp Asp Gly Asn Tyr Lys Thr Arg Ala Glu Val Lys Phe Glu Gly Asp Thr 420
Leu Val Asn Arg Ile Glu Leu Lys Gly Ile Asp Phe Lys Glu Asp Gly Asn Ile Leu Gly 440
His Lys Leu Glu Tyr Asn Tyr Ile Ser His Asn Val Tyr Ile Thr Ala Asp Lys Gln Lys 460
Asn Gly Ile Lys Ala Asn Phe Lys Ile Arg His Asn Ile Glu Asp Gly Ser Val Gln Leu 480
Ala Asp His Tyr Gln Gln Asn Thr Pro Ile Gly Asp Gly Pro Val Leu Leu Pro Asp Asn 500
His Tyr Leu Ser Thr Gln Ser Ala Leu Ser Lys Asp Pro Asn Glu Lys Arg Asp His Met 520
Val Leu Leu Glu Phe Val Thr Ala Ala Leu Glu His His His His His His End         538
```

**Figure S6.** Amino acid sequence of GR2-CFP. Colors correspond to the description in Figure S5.

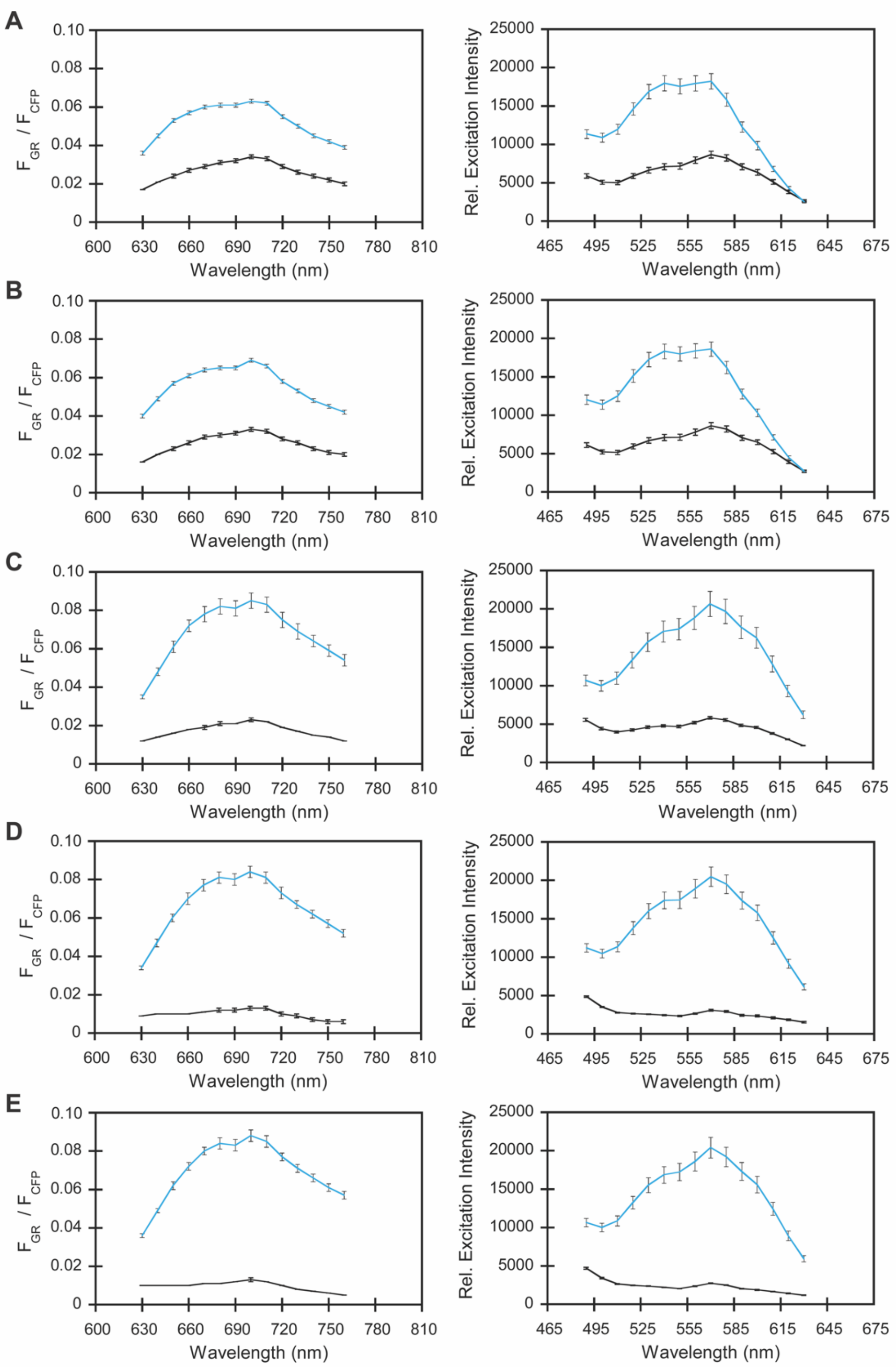


**Figure S7.** Characterization of the engineered GR1-CFP variants in *Escherichia coli* treated with 0 (black) and 400 mM sodium chloride (blue) in 50 mM sodium acetate buffer at pH 5. The emission spectrum of the rhodopsin normalized by the CFP emission at 485 nm (left) and excitation spectrum of the rhodopsin emission at 710 nm (left) is shown for (A) GR1-CFP, (B) GR1 E132K-CFP, (C) GR1 E132K/A84K-CFP, (D) GR1 E132K/A84K/T125C-CFP, and (E) GR2-CFP.

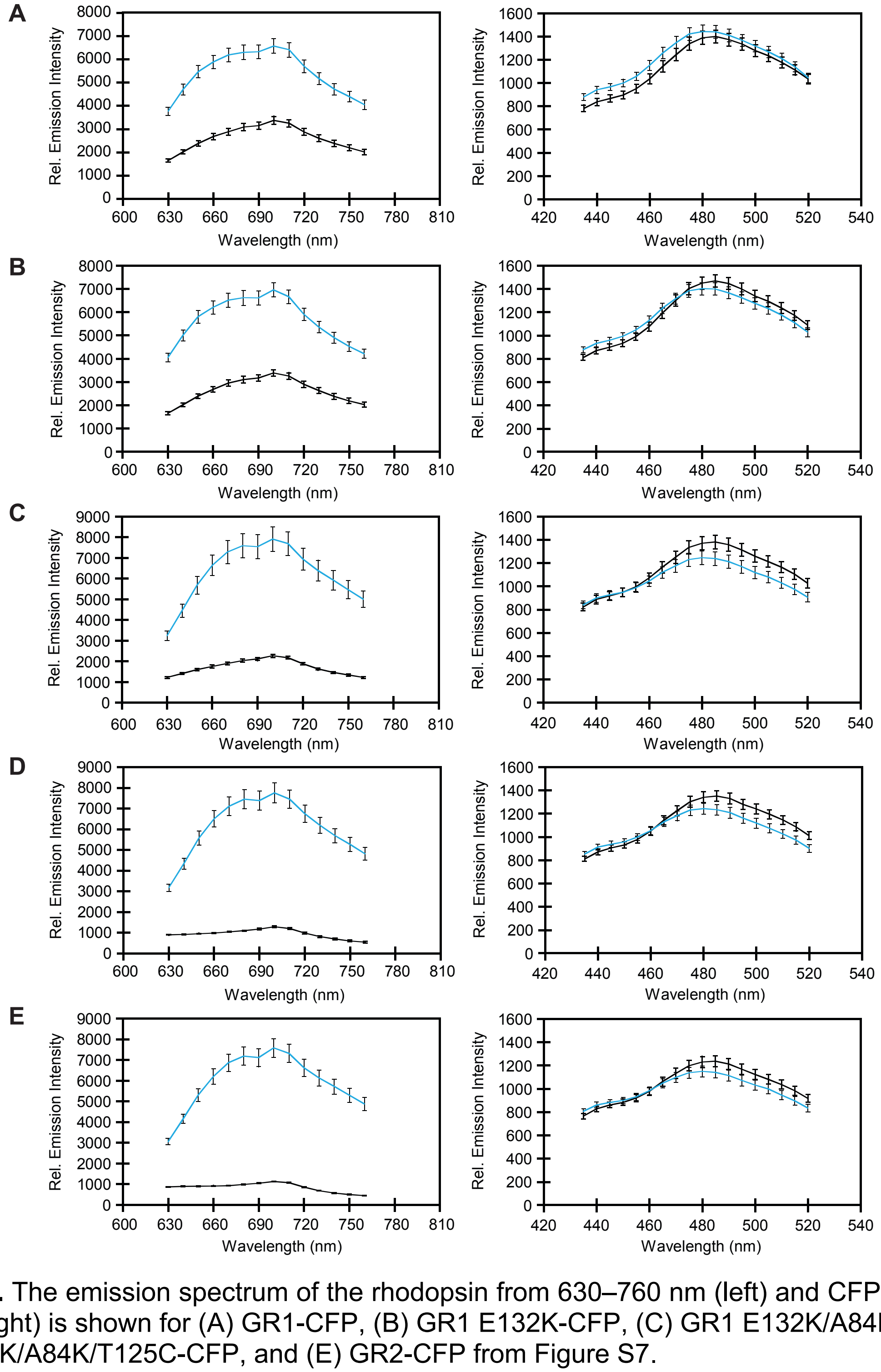


**Figure S8.** The emission spectrum of the rhodopsin from 630–760 nm (left) and CFP from 435–520 nm (right) is shown for (A) GR1-CFP, (B) GR1 E132K-CFP, (C) GR1 E132K/A84K-CFP, (D) GR1 E132K/A84K/T125C-CFP, and (E) GR2-CFP from Figure S7.

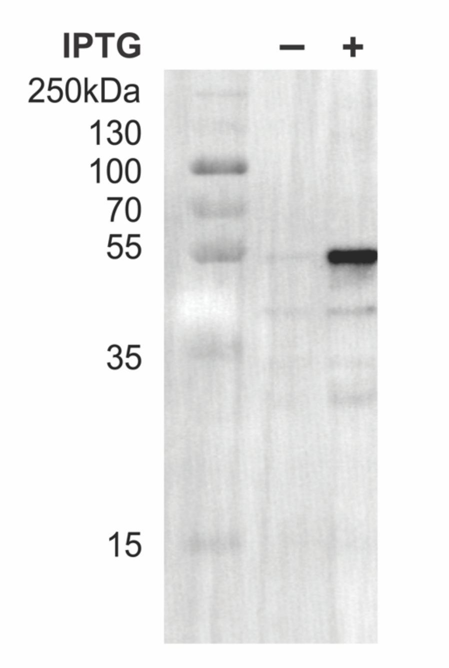


**Figure S9.** Representative western blot analysis using anti-6x-His tag antibody shows that GR2-CFP is expressed in the isopropyl β-D-1-thiogalactopyranoside (IPTG) induced sample. The expected molecular weight for the monomer is ~59.3 kDa.

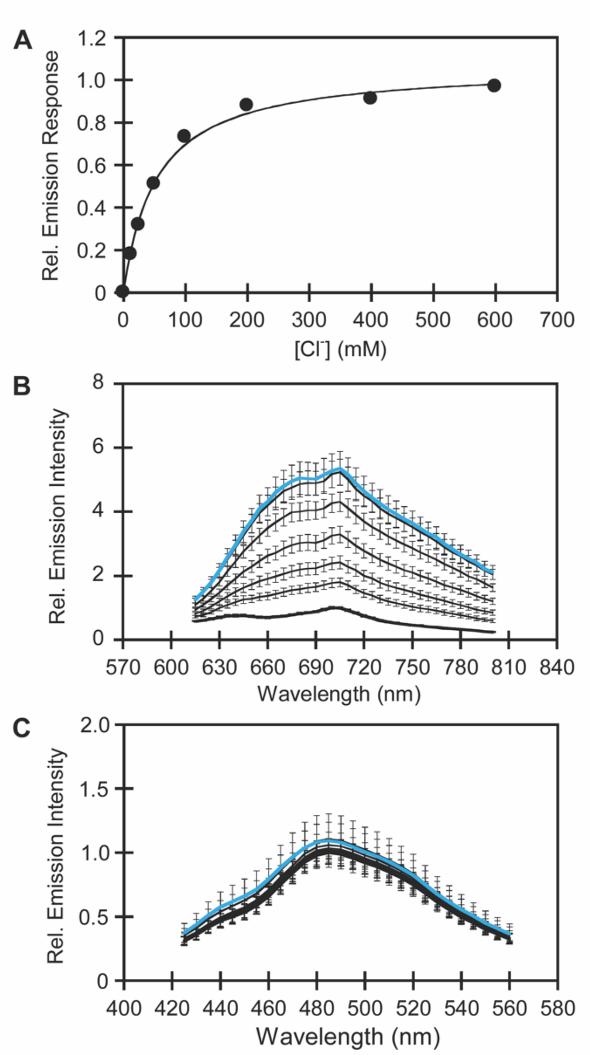


**Figure S10.** (A) Normalized integrated emission response ($F_{GR2}/F_{CFP}$) of GR2-CFP expressed in *E. coli* to 0, 12.5, 25, 50, 100, 200, 400, and 600 mM sodium chloride in 50 mM sodium acetate buffer at pH 5 to determine the apparent dissociation constant ($K_d$) from Figure 4A. Note: the additional data point for 600 mM sodium chloride is not shown in Figure 4A. The $K_d$ for $Cl^-$ is 53 ± 14 mM. (B) Emission spectra of GR2 for GR2-CFP from Figure 4A. (C) Emission spectra of CFP for GR2-CFP from Figure 4A. For the rhodopsin, the excitation was provided at 570 nm, and the emission was collected and integrated from 615–800 nm ($F_{GR2}$). For CFP, the excitation was provided at 390 nm, and the emission was collected from 425–560 nm. The emission spectra of GR2 at each point were normalized by the CFP emission intensity at 485 nm ($F_{CFP}$). The average of nine biological replicates from three technical replicates with standard error of the mean is shown.

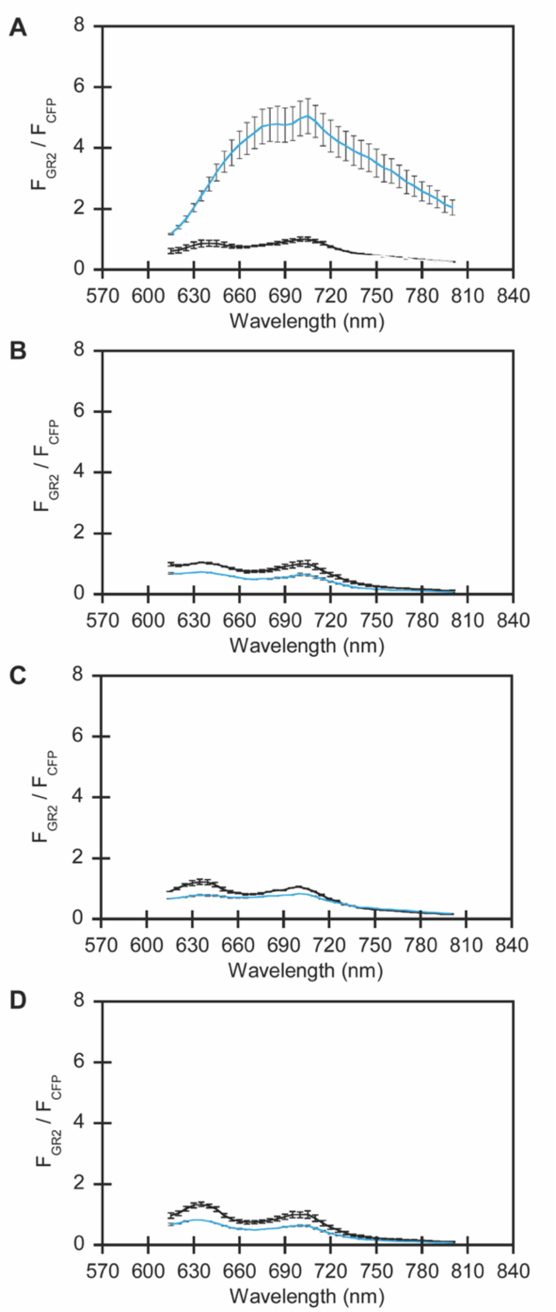


**Figure S11.** Normalized emission spectra of GR2-CFP in *E. coli* treated with 0 (black) or 400 mM (blue) sodium chloride in 50 mM sodium acetate buffer at pH 5 following expression (A) with IPTG and all-*trans*-retinal (ATR), (B) with IPTG only, (C) with ATR only, and (D) without IPTG and ATR. For the rhodopsin, the excitation was provided at 570 nm, and the emission was collected from 615–800 nm. For CFP, the excitation was provided at 390 nm, and the emission was collected from 425–560 nm. The emission spectra of GR2 at each point were normalized by the CFP emission intensity at 485 nm ($F_{CFP}$). The average of nine biological replicates from three technical replicates with standard error of the mean is shown.

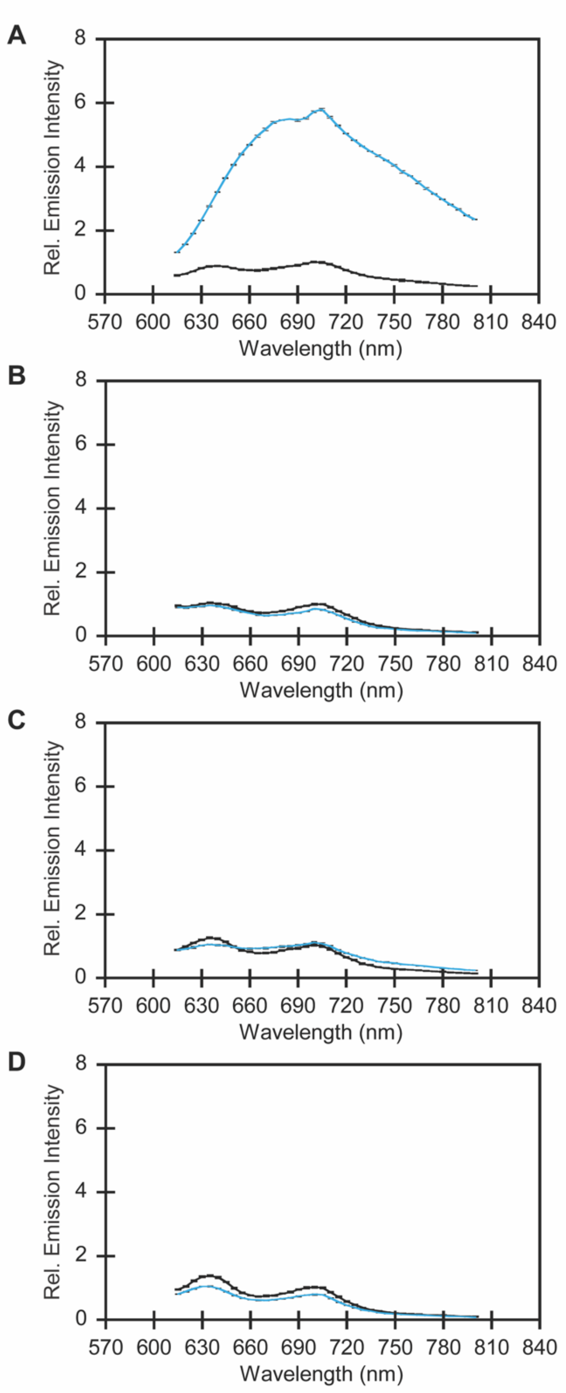


**Figure S12.** Emission spectra of GR2 for GR2-CFP in *E. coli* treated with 0 (black) or 400 mM (blue) sodium chloride following expression (A) with IPTG and ATR, (B) with IPTG only, (C) with ATR only, and (D) without IPTG and ATR from Figure S11. For the rhodopsin, the excitation was provided at 570 nm, and the emission was collected from 615–800 nm. The average of nine biological replicates from three technical replicates with standard error of the mean is shown.

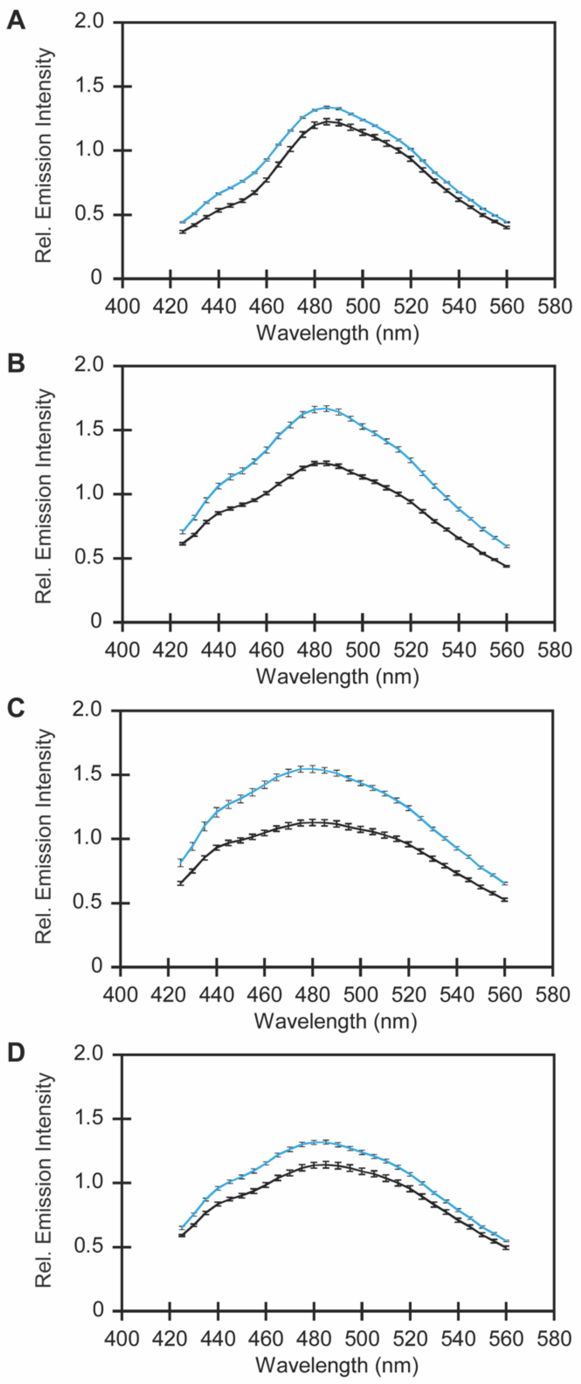


**Figure S13.** Emission spectra of CFP for GR2-CFP in *E. coli* treated with 0 (black) or 400 mM (blue) sodium chloride following expression (A) with IPTG and ATR, (B) with IPTG only, (C) with ATR only, and (D) without IPTG and ATR from Figure S11. For CFP, the excitation was provided at 390 nm, and the emission was collected from 425–560 nm. The average of nine biological replicates from three technical replicates with standard error of the mean is shown.

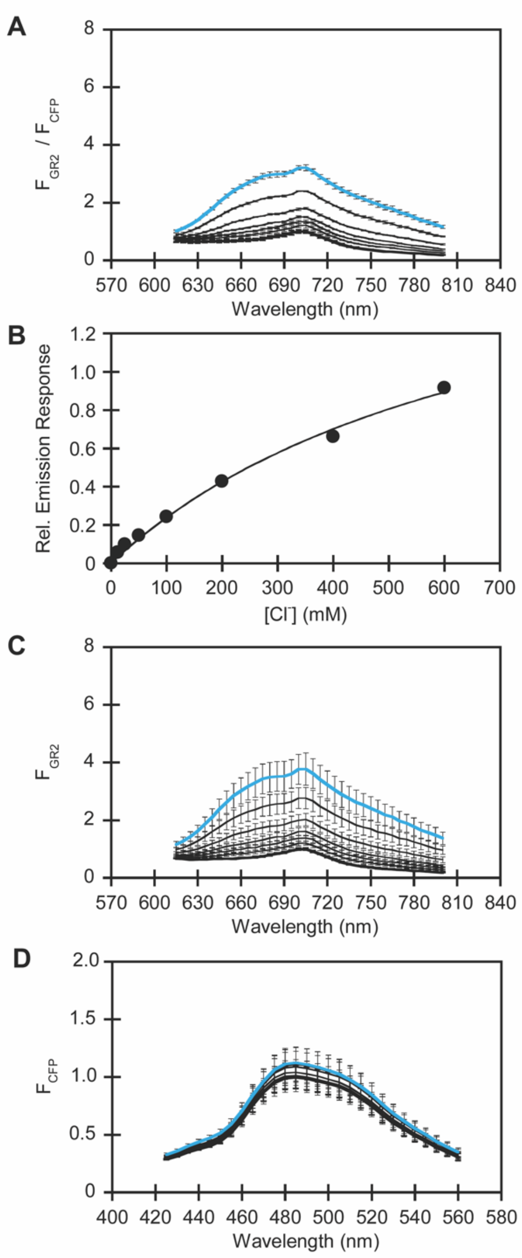


**Figure S14.** (A) Normalized emission spectra of GR2-CFP in *E. coli* treated with 0 (black and bold), 12.5, 25, 50, 100, 200, and 400 mM (blue) sodium chloride in 50 mM sodium phosphate buffer at pH 6. (B) Normalized integrated emission response ($F_{GR2}/F_{CFP}$) from *A* to determine the $K_d$. Note: the additional data point for 600 mM sodium chloride is not shown in *A*. The $K_d$ for $Cl^-$ is 731 ± 31 mM. (C) Emission spectra of GR2 from *A*. (D) Emission spectra of CFP from *A*. For the rhodopsin, the excitation was provided at 570 nm, and the emission intensity was collected and integrated from 615–800 nm ($F_{GR2}$). For CFP, the excitation was provided at 390 nm, and the emission was collected from 425–560 nm ($F_{CFP}$). The emission spectra of GR2 at each point were normalized by the CFP emission intensity at 485 nm ($F_{CFP}$). The average of nine biological replicates from three technical replicates with standard error of the mean is shown.

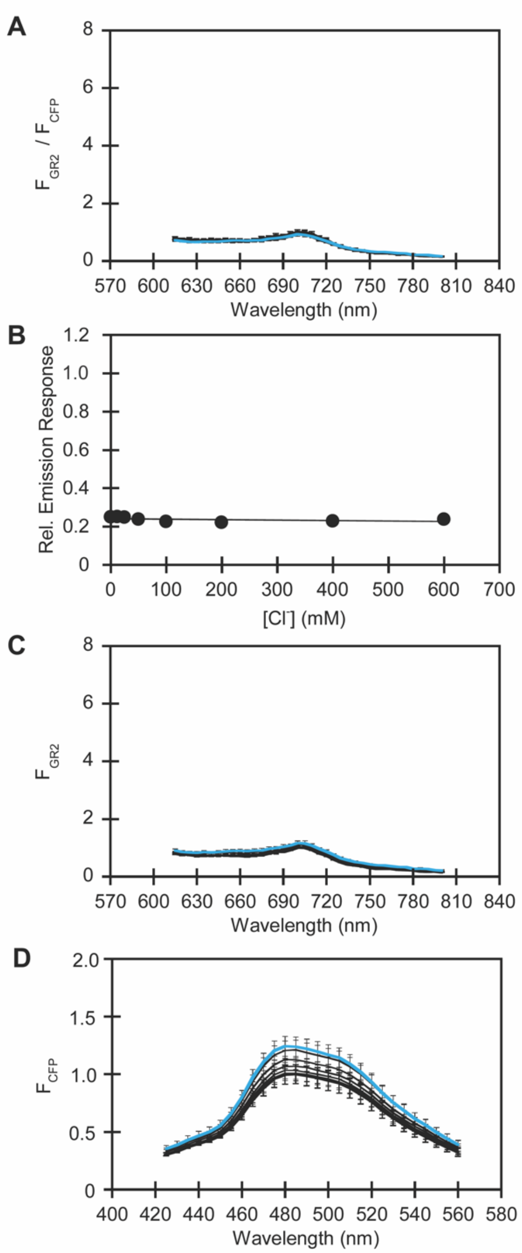


**Figure S15.** (A) Normalized emission spectra of GR2-CFP in *E. coli* treated with 0 (black and bold), 12.5, 25, 50, 100, 200, and 400 mM (blue) sodium chloride in 50 mM sodium phosphate buffer at pH 7. (B) Normalized integrated emission response ($F_{GR2}/F_{CFP}$) from *A.* Note: the additional data point for 600 mM sodium chloride is not shown in *A*. The $K_d$ could not be determined. (C) Emission spectra of GR2 from *A*. (D) Emission spectra of CFP from *A*. For the rhodopsin, the excitation was provided at 570 nm, and the emission intensity was collected and integrated from 615–800 nm ($F_{GR2}$). For CFP, the excitation was provided at 390 nm, and the emission was collected from 425–560 nm ($F_{CFP}$). The emission spectra of GR2 at each point were normalized by the CFP emission intensity at 485 nm ($F_{CFP}$). The average of nine biological replicates from three technical replicates with standard error of the mean is shown.

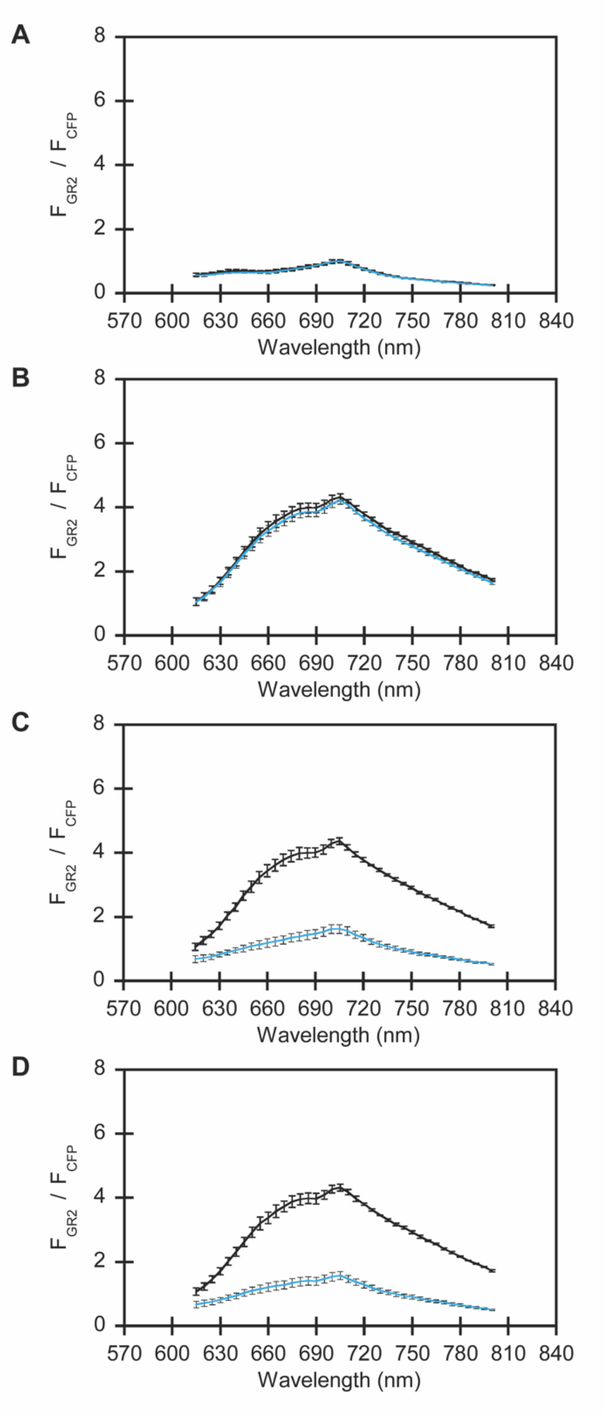


**Figure S16.** Normalized emission spectra of GR2-CFP in *E. coli* treated first in the presence of buffer only (control) or 400 mM sodium chloride (black) in 50 mM sodium acetate buffer at pH 5, followed by washing with (A) buffer only for the control sample, and (B) 400 mM sodium chloride, (C) buffer, or (D) 400 mM sodium gluconate for the sodium chloride treated samples (blue) for Figure 4B. For the rhodopsin, the excitation was provided at 570 nm, and the emission was collected from 615–800 nm. For CFP, the excitation was provided at 390 nm, and the emission was collected from 425–560 nm. The emission spectra of GR2 at each point were normalized by the CFP emission intensity at 485 nm ($F_{CFP}$). The average of nine biological replicates from three technical replicates with standard error of the mean is shown.

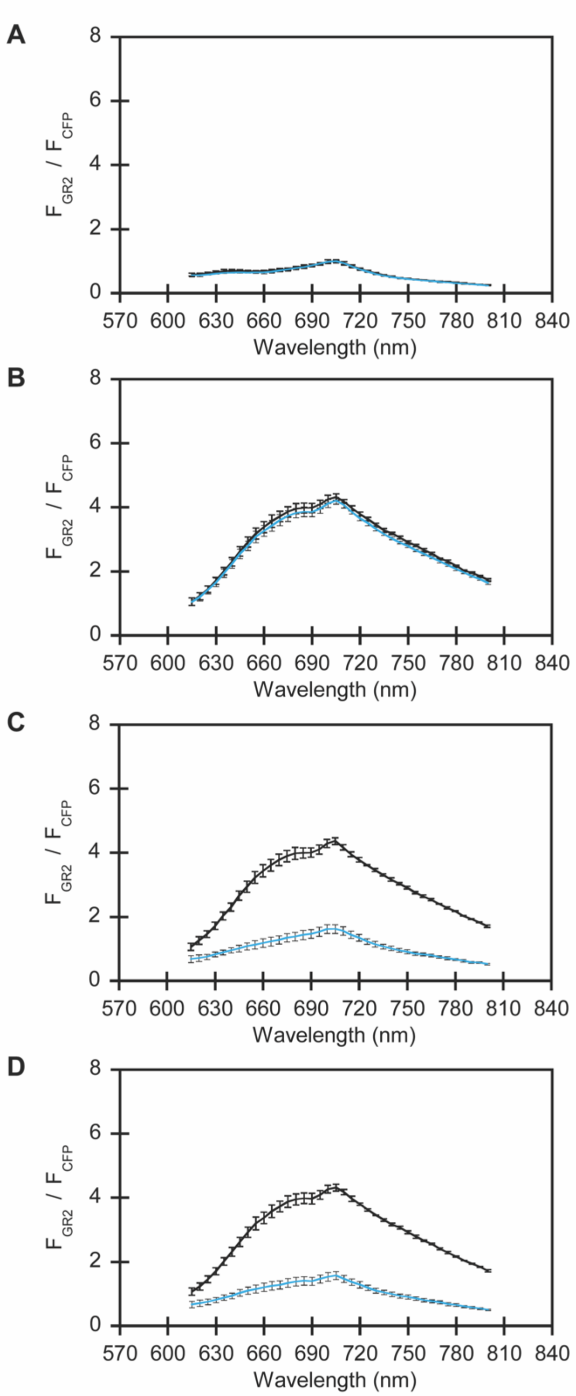


**Figure S17.** Emission spectra of GR2 for GR2-CFP in *E. coli* treated first in the presence of buffer only (control) or 400 mM sodium chloride (black), followed by washing with (A) buffer only for the control sample, and (B) 400 mM sodium chloride, (C) buffer, or (D) 400 mM sodium gluconate for the sodium chloride treated samples (blue) from Figure S16. For the rhodopsin, the excitation was provided at 570 nm, and the emission was collected from 615–800 nm. The average of nine biological replicates from three technical replicates with standard error of the mean is shown.

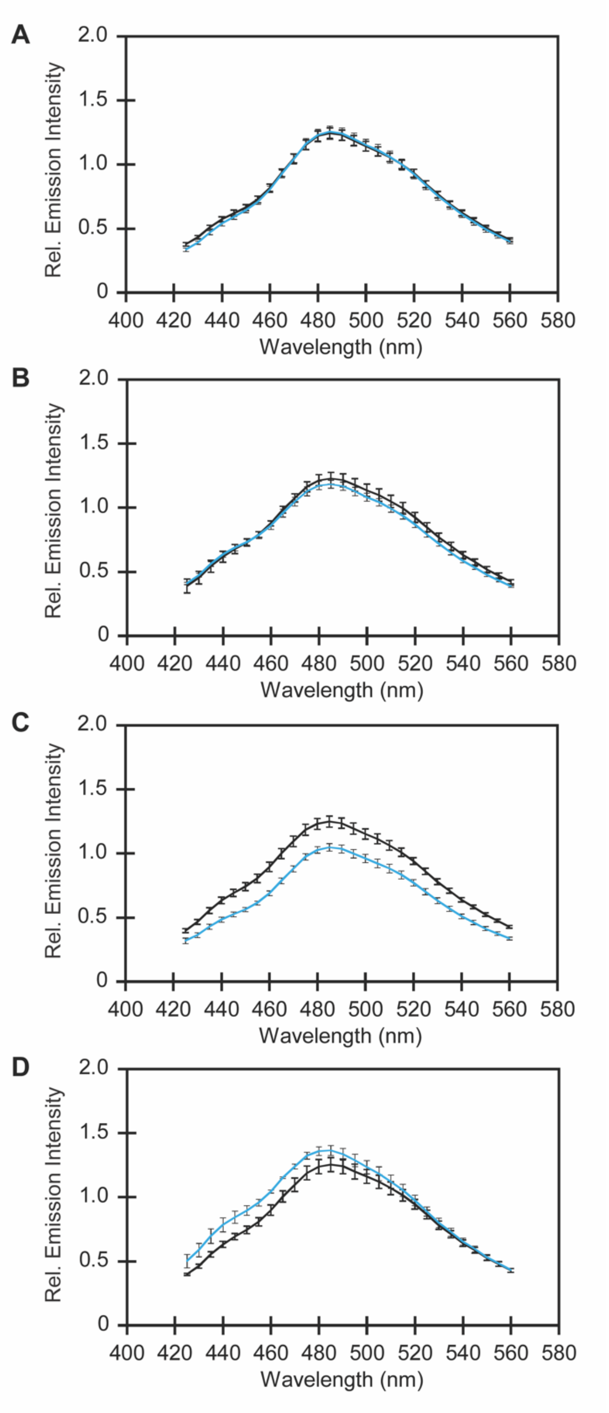


**Figure S18.** Emission spectra of CFP for GR2-CFP in *E. coli* treated first with buffer only (control) or 400 mM sodium chloride (black), followed by washing with (A) buffer only for the control sample, and (B) 400 mM sodium chloride, (C) buffer, or (D) 400 mM sodium gluconate for the sodium chloride treated samples (blue) from Figure S16. For CFP, the excitation was provided at 390 nm and the emission was collected from 425–560 nm. The average of nine biological replicates from three technical replicates with standard error of the mean is shown.

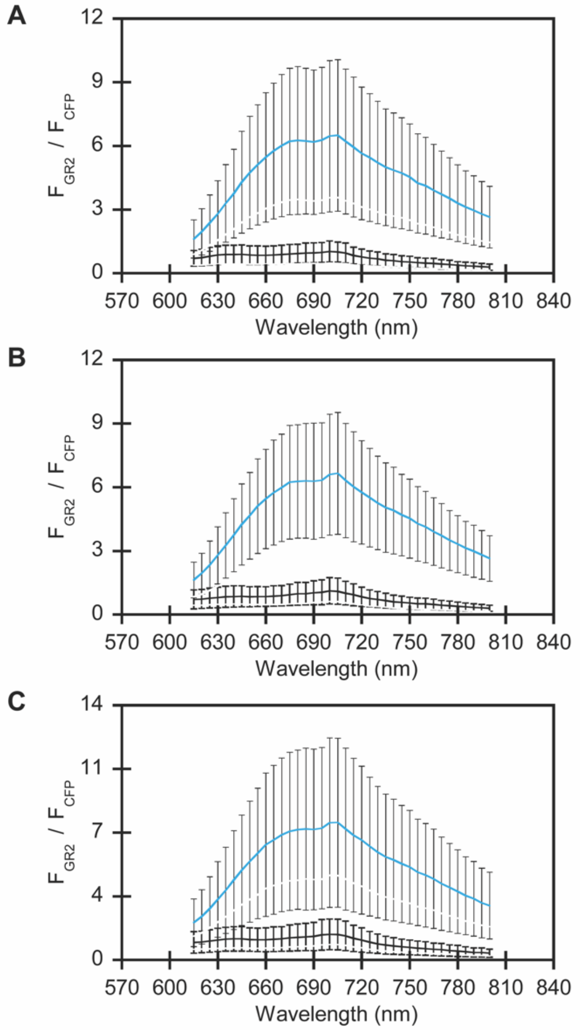


**Figure S19.** Normalized emission spectra of GR2-CFP in *E. coli* pre-treated with (A) 50 mM sodium acetate buffer at pH 5, (B) 0.3% DMSO, or (C) 30 μM CCCP with 0 (black) and 400 (blue) mM sodium chloride from Figure 4C. For the rhodopsin, the excitation was provided at 570 nm, and the emission was collected from 615–800 nm. For CFP, the excitation was provided at 390 nm, and the emission was collected from 425–560 nm. The emission spectra of GR2 at each point were normalized by the CFP emission intensity at 485 nm ($F_{CFP}$). The average of nine biological replicates from three technical replicates with standard error of the mean is shown.

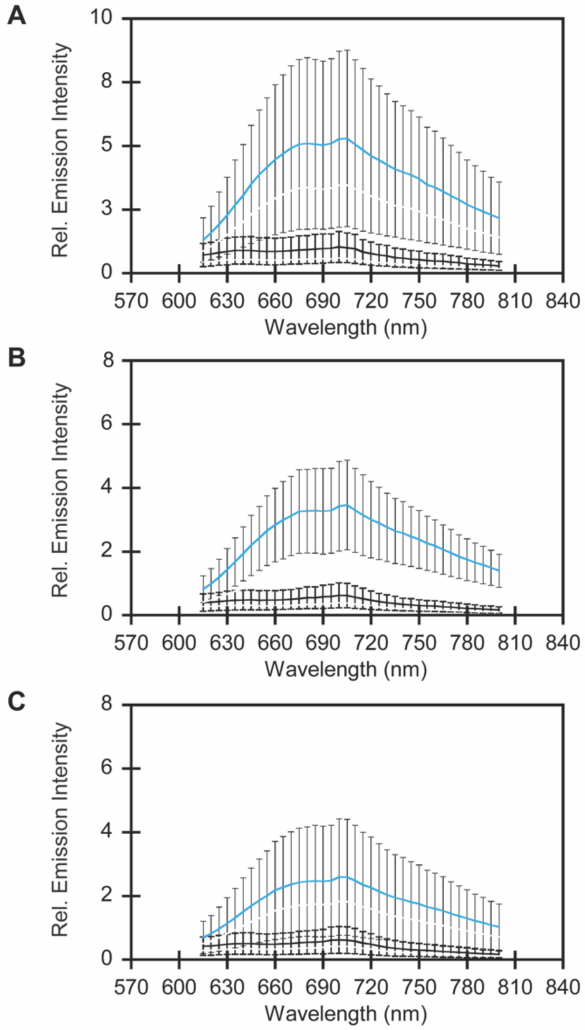


**Figure S20.** Emission spectra of GR2 for GR2-CFP in *E. coli* pre-treated with (A) 50 mM sodium acetate buffer at pH 5, (B) 0.3% DMSO, or (C) 30 μM CCCP with 0 (black) and 400 (blue) mM sodium chloride from Figure S19. For the rhodopsin, the excitation was provided at 570 nm, and the emission was collected from 615–800 nm. The average of nine biological replicates from three technical replicates with standard error of the mean is shown.

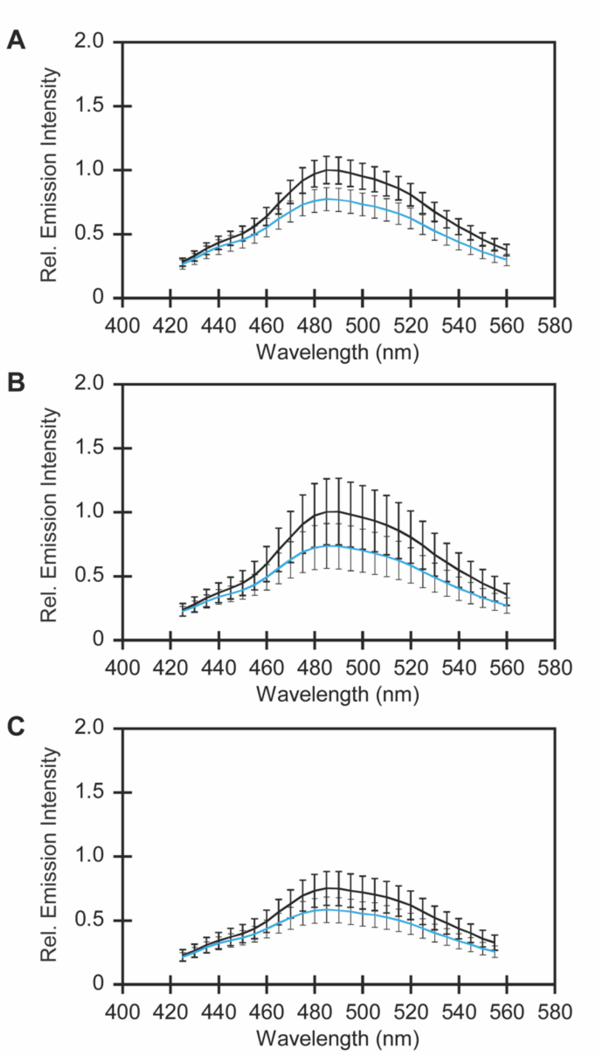


**Figure S21.** Emission spectra of CFP for GR2-CFP in *E. coli* pre-treated with (A) 50 mM sodium acetate buffer at pH 5, (B) 0.3% DMSO or (C) 30 µM CCCP with 0 (black) and 400 (blue) mM sodium chloride from Figure S19. For CFP, the excitation was provided at 390 nm, and the emission was collected from 425–560 nm. The average of nine biological replicates from three technical replicates with standard error of the mean is shown.

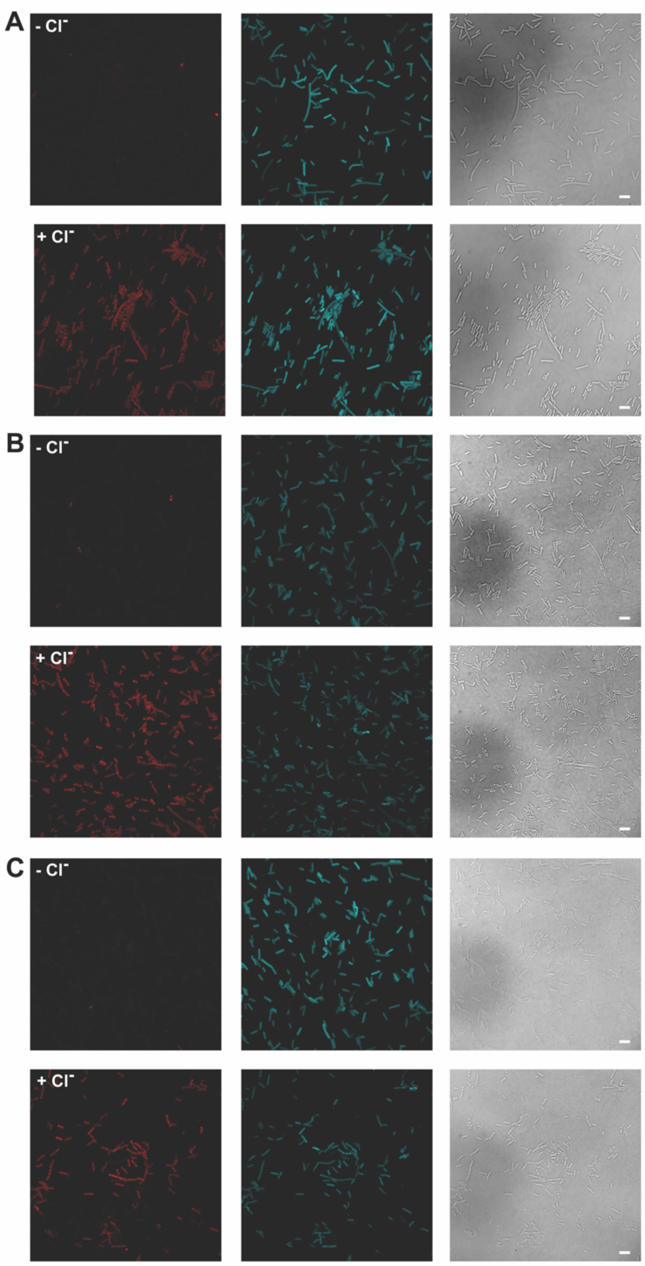


**Figure S22.** Representative confocal fluorescence microscopy images from three different biological replicates (A–C) are shown for *E. coli* expressing GR2-CFP on agarose pads with 0 mM and 400 mM sodium chloride in 50 mM acetate buffer at pH 5. At least five fields were imaged for each biological replicate. In each panel, the GR2 emission (red) is on the left, the CFP emission (cyan) is in the middle, and the differential interference contrast (DIC) image is on the right (scale bar = 5 μm). Analysis is shown in Figure 5 and S23.

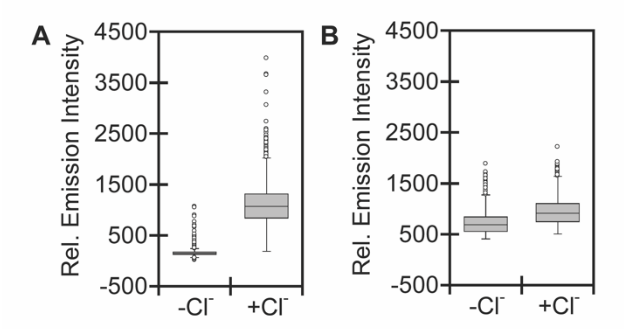


**Figure S23.** Boxplots show the emission response of (A) GR2 and (B) CFP in each cell analyzed from three biological replicates (n = 2,583 regions of interest (ROIs) for 0 mM sodium chloride; n = 2,367 ROIs for 400 mM sodium chloride). The gray boxes correspond to the lower and upper quartile data with the minimum and maximum values extending below and above the box. The median values are indicated by the black lines in the gray boxes, and outliers are shown as open circles.

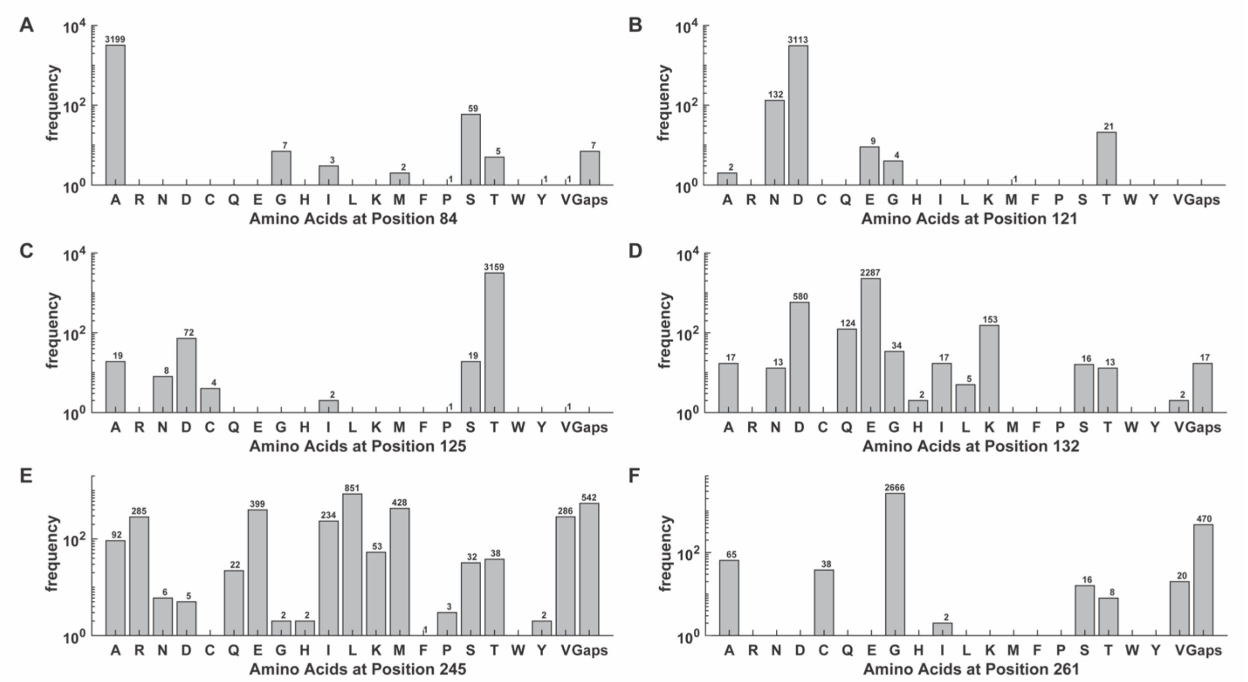


**Figure S24.** Composition of amino acids at each position along the proton-pumping pathway for all proteins in the rhodopsin family. The amino acid is shown on the x-axis with the number of proteins on the y-axis. If there is no amino acid at a given position in the alignment, it is defined as a gap.
